# Enabling subcellular DESI-MSI for broad adoption: Acquisition, analysis, and application

**DOI:** 10.64898/2026.09.25.754519

**Authors:** Daniel J. Boehmler, Melanie Loth, Tristan O’Harrow, Giang Hoang, Om B. Patel, Farheen Akhtar, Pinky Kain, Arjun Sengupta, Georgios Paschos, Mingyao Li, Terence P. Gade, Garret A. FitzGerald, Aalim M. Weljie

**Author notes:** **Corresponding Author:** Aalim M. Weljie, Ph.D., 10-196 Smilow Center for Translational Research, 3400 Civic Center Blvd, Philadelphia PA 19104.

## Abstract

Subcellular mass spectrometry imaging (MSI) has required specialized instrumentation and approaches. Additionally, data analysis has required expensive proprietary software, or code-based open-source tools. These barriers render spatial metabolomics less accessible than peer spatial omics techniques (transcriptomics, proteomics). Here we demonstrate subcellular MSI on standard commercial hardware, analyzed using code-free, open-source software. An inexpensive, fully reversible modification of a stock DESI source takes the platform past its typical 5-10 µm pixels to routine 2×2 µm pixels, and 1×1 µm in proof-of-principle experiments. At a tissue-glass boundary imaged at 2×2 µm, tissue-specific ions showed no measurable spillover onto the adjacent glass within the ±4.4 µm uncertainty of the edge position. To make these data interpretable, we extend MSI.EAGLE, an open-source, vendor-agnostic application with two new methods: i) Spatial UMAP preserves tissue architecture alongside subcellular detail, where conventional clustering breaks apart at high resolution; ii) Histology Fit performs landmark-free co-registration, and recovers synthetic misalignments of up to 8 µm to within one pixel of a reference registration. Together, these enable previously high-burden spatial metabolomics experiments. In mouse brain tissue, we demonstrate genuine subcellular signal by resolving mitochondrial cardiolipin into perinuclear puncta, quantifying cytoplasmic enrichment across 6,073 cells. Cell identities are assigned to MSI pixels using histology-predicted transcriptomics, linking cell provenance to metabolism in 2D space directly. Finally, we recover cell-type-specific metabolomes consistent with the transcriptomic identities. This workflow allows cellular and subcellular spatial metabolomics linked to cell provenance, with only stock MSI hardware, open-source GUI-based software, and common histology techniques, putting it within reach of many researchers and laboratories.

## Intro

Spatial biology connects molecular profiling of proteins^1,2^, transcripts^3,4^, and metabolites^5^ to visual, anatomical data. Traditional bulk methods forgo this spatial context, and are thus limited in their ability to explain the influence of tissue structure, neighborhood interactions, and microenvironments on cell behavior and disease states.

Advances in the acquisition and analysis of spatial proteomics and spatial transcriptomics data have been transformative, leading to their wide adoption by researchers. In pursuit of high-fidelity, reproducible results, the standardization of sample preparation, data acquisition, and analysis pipelines are becoming more common for these techniques. However, the same cannot be said for spatial metabolomics.

Spatial metabolomics workflows vary widely; Sample slide preparation, ionization method, hardware modifications, data formats, and analysis platforms are often tailored to each laboratory’s needs. While this kind of specialization lends itself to high-quality data, it simultaneously limits accessibility to the broader research community. The bespoke nature of spatial metabolomics is particularly evident in high spatial resolution mass spectrometry imaging (MSI), where analysis at the cellular and subcellular level is 1) difficult to interpret, 2) computationally intensive, and 3) often requires specific, expensive hardware additions for acquisition.

Vendor-supplied MSI instrumentation has steadily advanced year-over-year, with improvements in both spatially-enabled ionization source hardware and mass spectrometer quality increasing spatial resolution and sensitivity, respectively. Matrix-assisted Laser Desorption Ionization (MALDI) and Desorption Electrospray Ionization (DESI) are currently the first and second most widely utilized ionization techniques, respectively.^6^ Both platforms provide rich molecular coverage, relative ease-of-use, and the ability to acquire diverse sample types and sizes, meaning they have the most generalized utility to researchers versus more specialized techniques. DESI is an ambient technique that does not require matrix application (like MALDI) to assist in ionization, conferring several advantages. The lack of a chemical matrix removes a sample preparation step that could introduce variation and requires extra equipment, avoids the need for matrix removal prior to post-acquisition secondary assays and stains, and means DESI is truly label-free, working on tissues in their native state. MALDI has a clear edge in resolution: with MALDI-2 - a configuration requiring the addition of a secondary post-ionization laser - pixel sizes of 1×1 to 2×2 µm are achievable.^7^ Conventional DESI currently achieves ∼5 µm pixels by i) optimizing spray parameters^8^ and ii) adding tubing and a column to the solvent flow path to raise pump backpressure and stabilize the spray, both simple and inexpensive hardware changes.^9^

Advancement of analysis pipelines for this kind of high-resolution MSI lags behind the described hardware advances, particularly rigorous co-registration with secondary techniques at the cellular and sub-cellular scale. MSI analyses are commonly performed with SCiLS Lab by Bruker on the commercial end, and the Cardinal package in R as an open-source solution.^10^ However, the ideal for most researchers is a unified, open-source, MSI vendor-agnostic, code-free analysis layer with landmark-free secondary assay co-registration, built in statistical workflows, and high-quality figure generation. Current tools address these points individually, but few allow beginning-to-end analysis in one code-free interface.

Here, we present the development of hardware and software solutions to democratize the acquisition and analysis of cellular and sub-cellular MSI. Our approach involves 1) acquisition of cellular and sub-cellular MSI data at 1 × 1 and 2×2 µm using a minimally adapted DESI-XS source, and 2) end-to-end analysis with our open-source software package, MSI.EAGLE.^11^ Starting with a mass spectrometry image (MALDI or DESI) in .imzML format, an H&E stain of the same tissue, and consumer-grade compute, a researcher can obtain cellular and sub-cellular metabolic information linked to cell provenance with our workflow. The entire analysis can be completed without difficult or expensive hardware modification, without writing a single line of code, and using only open-source software. By lowering the barrier to entry in this manner, we hope to bring the state of high-resolution spatial metabolomics on par with other spatial omics techniques in its accessibility to researchers.

## Results

### Subcellular mass spectrometry imaging with a commercial DESI source

Acquiring mass spectrometry imaging data at the cellular and sub-cellular scale is often the first barrier researchers encounter. DESI-MSI is a more accessible platform due to the aforementioned sample preparation advantages, and the cost being less than half that of an equivalent MALDI platform.^12,13^ Thus, we endeavored to resolve the gap in pixel size capability between the two platforms.

We began by extending the low-flow DESI configuration of Towers et al.^9^ on a Waters DESI-XS source coupled to a Xevo G2-XS QTOF. In short, we added two in-line C18 columns and 30 meters of fused-silica capillary tubing along the solvent flow path and decreased the emitter tip-to-sample distance. A full description of these hardware modifications is provided in the methods. This simple, reversible, and inexpensive modification raised backpressure beyond the original configuration, tightening the spray footprint (**Fig. 1a, Supplementary Fig. 1)**. With this setup we acquired data at both 2×2 µm and, in proof-of-principle experiments, 1×1 µm pixel sizes.

**Fig. 1.**
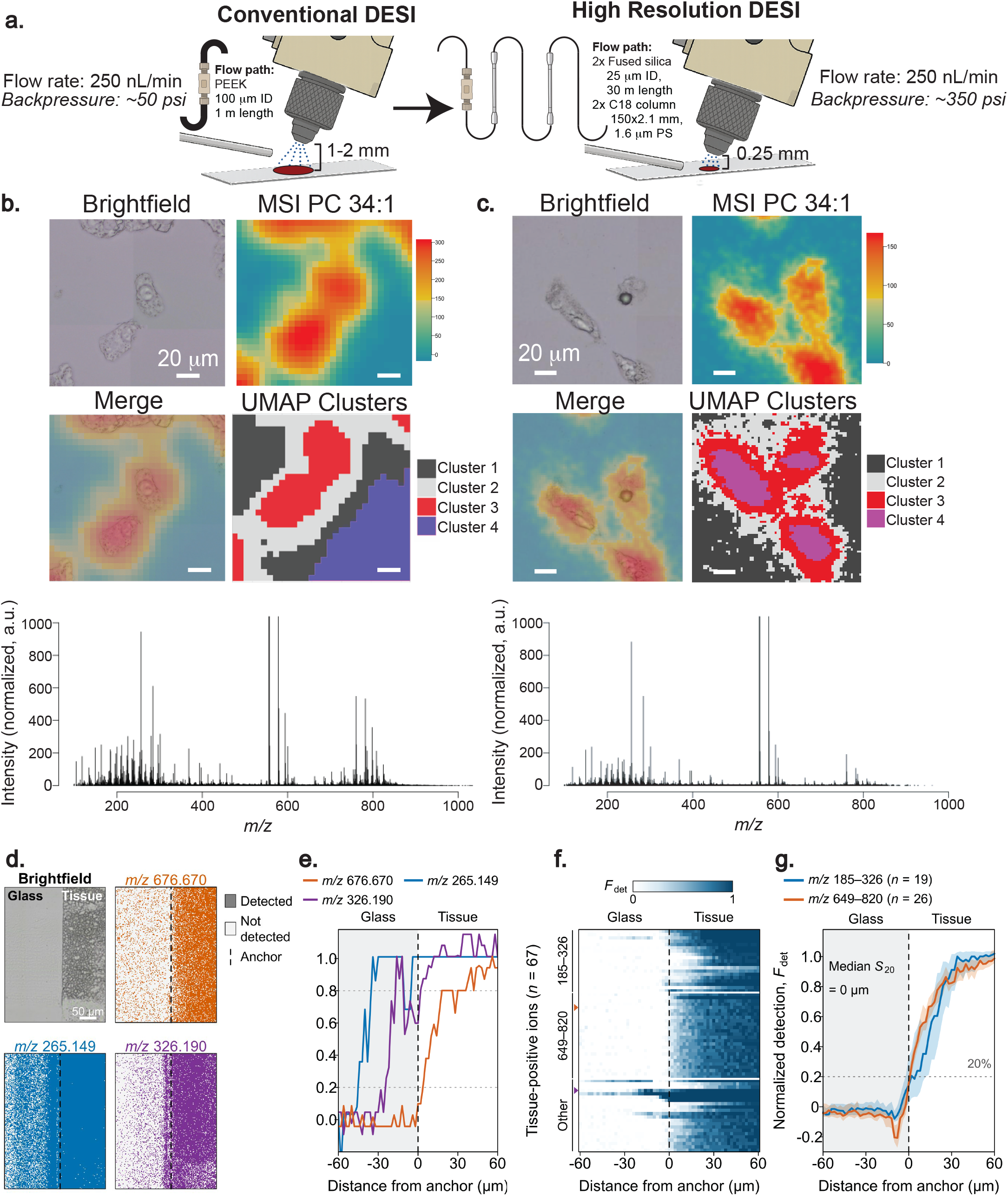
Subcellular DESI-MSI with a minimally modified commercial source. **a,** Solvent-flow-path modification converting a conventional DESI-XS source (PEEK line, 100 µm i.d., 1 m; emitter-to-sample ∼1-2 mm; ∼50 psi) to the high-resolution configuration (2x in-line C18 columns, 150×2.1 mm, 1.6 µm; 2x fused-silica capillary, 25 µm i.d., 30 m; emitter-to-sample 0.25 mm; ∼350 psi) at a constant 250 nL min^-1^ spray. **b,c,** Positive-mode acquisition of SNU-449 hepatocellular carcinoma cells cultured on glass at 5×5 µm (**b**) and 2×2 µm (**c**), each showing the brightfield image, the PC 34:1 ion image, the brightfield-ion merge, and the Spatial UMAP cluster map (four clusters), with the mean mass spectrum below. Cell outlines sharpen and intracellular clusters (e.g., Cluster 4) appear only at 2×2 µm. Scale bars, 20 µm. **d-g,** Tissue-to-glass spillover at a mouse brain section edge imaged at 2×2 µm; glass is left and tissue right in all spatial panels. Distance is measured from a molecular anchor, the detection onset of a tissue-specific feature, where d = 0; negative values are on glass. **d,** Post-scan brightfield of the analyzed field (201×150 pixels) and per-pixel detection maps (colored, ion detected; pale, not detected) for a tissue-positive ion (*m/z* 676.670), an ion previously designated glass-enriched by abundance (*m/z* 265.149; its detection is nonetheless higher in tissue), and an interface-enriched ion (*m/z* 326.190). The dashed line marks the anchor. **e,** Normalized detection fraction (F_det_; far-glass baseline = 0, tissue = 1) for the ions in **d,** taken as the median across 20 ten-column blocks. Dotted lines mark 20% and 80%. The three ions change at different positions and with different shapes across the same edge, so a conventional 20-80% transition width mixes molecular localization, interface chemistry and spatial spreading, and is not read as spatial resolution. **f,** Median F_det_ of all 67 tissue-positive ions (after isotope/related-feature collapse), grouped as *m/z* 185-326 (n = 19), *m/z* 649-820 (n = 26) and other masses (n = 22), ordered by *m/z* within each group. Color is clipped to 0-1 for display only. Triangles mark *m/z* 676.670 (orange) and *m/z* 326.190 (purple). **g,** Group-median F_det_ for the two prespecified mass groups; ribbons are pointwise 95% intervals from 2,000 hierarchical block-bootstrap replicates. S₂₀ is the distance onto glass at which tissue-specific detection last reaches 20%; it is aggregated across block and ion values rather than read from the group curve. Median S₂₀ was 0 µm for both groups (within-section intervals 0-0.94 µm for *m/z* 185-326 and 0-0.30 µm for *m/z* 649-820; separate anchor uncertainty ±4.4 µm). S₂₀ is an upper bound on spillover, from one section. Scale bars: 20 µm (b,c); 50 µm (**d**).

To benchmark pixel size against a sample of known geometry, we imaged SNU-449 hepatocellular carcinoma cells cultured directly on glass slides and compared positive mode 5×5 µm and 2×2 µm acquisitions head-to-head (**Fig. 1b,c**). SNU-449 cells are 20-30 µm in diameter, comparable to many mammalian cells (typically 10-30 µm). A 2×2 µm pixel covers 4 µm² compared with 25 µm² for a 5×5 µm pixel, a ∼6-fold improvement in sampling area. Visual inspection with matched contrast and smoothing showed clearly sharper cell outlines at the smaller pixel size (**Fig. 1b,c**). Unsupervised clustering of the data demonstrated a quantitative improvement at 2×2 µm pixels; We projected spatial UMAP clusters (See **Fig. 3** and Methods) back onto the ion image. At 5 µm, cluster boundaries tracked individual cell outlines, representing the average metabolic signature of an entire cell. At 2 µm, multiple distinct clusters appeared within individual cells, revealing spatially organized intracellular chemical compartments (**Fig. 1c**).

A smaller pixel only improves spatial fidelity if the desorption spray does not carry analyte laterally beyond the sampled spot. To test this directly, we imaged the boundary between a mouse brain section and the bare glass beside it at 2×2 µm in negative ionization mode (**Fig. 1d**). Any tissue-derived signal on the glass side of this boundary would reveal redistribution by the spray or by sectioning. Because detection fraction does not depend on total-ion-current normalization, we used it as the readout: the fraction of pixels in which an ion is detected, measured in 20 ten-pixel-wide blocks along the edge. We measured distance from a molecular anchor, the detection onset of a tissue-specific feature, rather than from the optical edge, which cannot be registered precisely at this scale. Individual ions did not behave alike at the boundary (**Fig. 1d,e**). A tissue-positive ion (*m/z* 676.670) remained at glass baseline up to the anchor and reached 20% of its tissue detection level only 3.7 µm into the tissue. By contrast, an ion classed as glass-enriched by abundance (*m/z* 265.149) and an interface-enriched ion (*m/z* 326.190) changed 20-45 µm before the anchor, the latter with a local maximum at the interface. Because these transitions differ in position and shape across the same edge, a conventional 20-80% edge width mixes molecular localization and interface chemistry with any physical spreading, and we do not interpret it as spatial resolution. We therefore used a one-sided measure: S₂₀, the distance onto glass at which tissue-specific detection last reaches 20%. Across all 67 tissue-positive ions (**Fig. 1f**), glass-side detection was essentially absent. In both prespecified mass groups, detection stayed at the far-glass baseline up to the anchor and rose only within the tissue (**Fig. 1g**). Median S₂₀ was 0 µm for both low-mass (*m/z* 185-326; n = 19; within-section interval 0-0.94 µm) and higher-mass ions (*m/z* 649-820; n = 26; 0-0.30 µm), with a separate anchor uncertainty of ±4.4 µm. Thus, at 2×2 µm, tissue-specific signal did not extend onto the adjacent glass beyond the molecular edge by a measurable amount.

### Subcellular localization of mitochondrial cardiolipin in mouse hippocampus

MSI of tissue sections generally confer the most utility for researchers. Thus, we next acquired DESI-MSI in negative mode from the hippocampal region of a mouse brain at both 5×5 µm and 2×2 µm pixel sizes, with H&E staining performed on the same section after acquisition (**Fig. 2, Supplementary Table 1**). We chose to focus on hippocampal anatomy because it provides a natural test case for resolution, with well-characterized layers and cell types. For demonstrative purposes, we’ll focus on three distinct regions: The pyramidal layer, stratum oriens, and alveus.

**Fig. 2.**
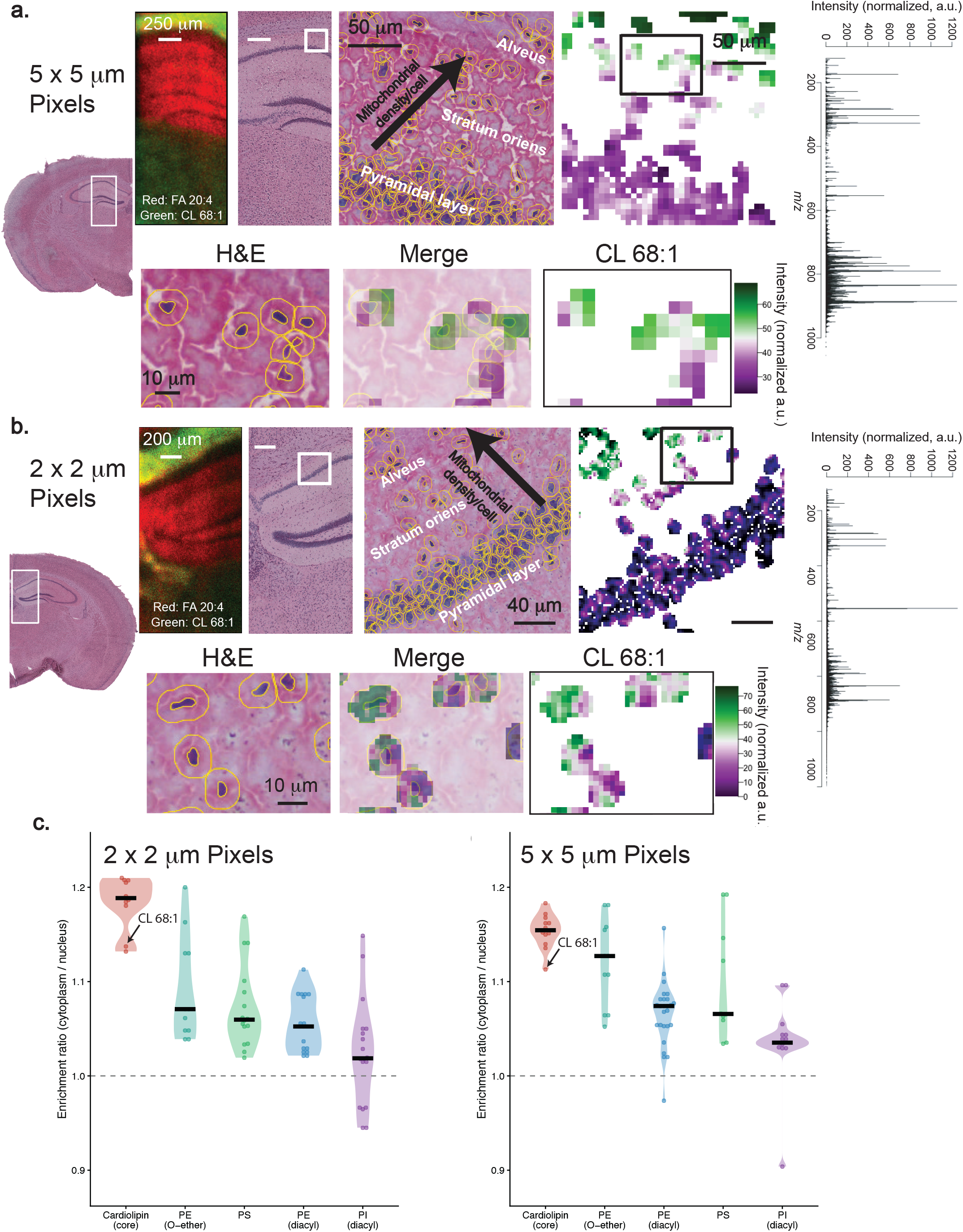
2×2 µm DESI-MSI localizes mitochondrial cardiolipin to the cytoplasm of the mouse hippocampus. Negative-mode DESI-MSI of the hippocampal region with post-acquisition H&E of the same section. **a,b,** Separate acquisitions at 5×5 µm (**a**) and 2×2 µm (**b**), each showing (left to right) a two-color overlay of FA 20:4 (red) and CL 68:1 (green), the H&E section with the imaged ROI boxed, a segmented H&E zoom (yellow cell outlines) spanning the pyramidal layer, stratum oriens and alveus (arrow, increasing mitochondrial density per cell), the CL 68:1 ion image, and the mean mass spectrum; the boxed region is enlarged below as H&E, merge and CL 68:1. CL 68:1 is uniform across each soma at 5×5 µm but resolves into perinuclear puncta at 2×2 µm. **c,** Cytoplasm-to-nucleus enrichment ratios at 2×2 µm (left) and 5×5 µm (right) for cardiolipin, PE (O-ether), PS, PE (diacyl) and PI (diacyl); each point is a compound, black bars are class medians, and the dashed line marks a ratio of 1. Cardiolipins (including CL 68:1, arrow) are the most cytoplasm-enriched class. Scale bars: overlays 250 µm (**a**) and 200 µm (**b**); segmented zooms 50 µm (**a**) and 40 µm (**b**); enlarged insets 10 µm.

At 5×5 µm pixel size, MS images reveal a gradient across these layers of increasing CL 68:1 abundance - a cardiolipin species which serves as a specific marker for the mitochondrial inner membrane (**Fig. 2a)**.^14^ This cardiolipin gradient is consistent with what we know about mitochondrial density in the soma of these cell types in young adult mice; Pyramidal neurons fire at lower sustained rates and distribute most of their mitochondria into dendrites and axons rather than concentrating them in the soma.^15^ The stratum oriens is dominated by fast-spiking inhibitory interneurons with intense metabolic demands and denser somatic mitochondrial populations.^16^ Oligodendrocytes in the alveus are among the most metabolically active cells in the brain, responsible for the high energetic cost of myelination, and so have the highest mitochondrial density.^17^

While highly detailed sub-organ and even cellular information can be gleaned at 5×5 µm pixel size, observation of the zoomed inset in **Fig. 2a** shows that signal is distributed uniformly across each cell body - there is no way to distinguish intracellular localization from cell-to-cell variation in total abundance. At 2×2 µm pixel size, the signal resolves into distinct puncta within individual cells, concentrated in the cytoplasmic compartment surrounding the nucleus where mitochondria (and thus, cardiolipins) localize **(Fig 2b)**. This pattern is not accessible at the coarser pixel size **(Fig 2a)**, where signal averaging across the cell area obscures it.

As further evidence of detectable subcellular localization in 2×2 µm data, we sought to compare the metabolism between pixels mapped to the cytoplasm versus pixels mapped to the nucleus in all 6073 cells contained in the dataset. We measured the abundance of 73 compounds in these pixel masks, then plotted the ratio between the cytoplasm and nucleus **(Fig. 2c)**. These perinuclear enrichment ratios show that our panel of mitochondrial cardiolipins are consistently the most enriched compound class in cytoplasmic annuli across 6073 cells: this panel of 10 high-confidence CL species was enriched in the cytoplasm by a mean of 18% (range 13-21%), versus only 8% (PE), 7% (PS) and 3% (PI) for the other major phospholipid classes; This is a 2.4-, 2.5- and 7.3-fold greater enrichment, respectively, indicating capture of genuine subcellular structure. Within the other three classes shown (PS, PE, PI), compounds that are ether-linked and/or less polyunsaturated than their class average are also enriched in the cytoplasm, consistent with their known mitochondrial association.^18,19^ On the other hand, highly polyunsaturated species (particularly those containing arachidonic acid, 20:4) trend toward nuclear localization, consistent with their known role in nuclear signaling.^20^ These patterns are also more prominent in 2×2 µm relative to 5×5 µm data (**Fig. 2c**): at 5×5 µm this contrast was compressed, with the cardiolipin panel enriched by only 15% while PS, PE and PI rose to 10%, 9% and 4%, reducing cardiolipin’s enrichment advantage to 1.6-, 1.8- and 4.2-fold, respectively. This indicates that 2×2 µm DESI-MSI provides a higher fidelity readout of mitochondrial membrane lipid distribution at the subcellular level.

### MSI.EAGLE: an integrated platform for end-to-end MSI analysis

To enable robust and facile analysis of data acquired with both our method and others, we developed MSI.EAGLE, an open-source R Shiny application that provides an end-to-end workflow from raw.imzML import, histology image co-registration, and publication-quality figure export, without requiring programming expertise (**Supplementary Fig. 2**). MSI.EAGLE is built on the Cardinal framework, and contains 9 distinct analysis modules, with each including an extensive set of advanced processing features that. Novel in this release are Spatial UMAP and Histology Fit workflows, which improve unsupervised clustering of high-resolution MSI data and enable quantitative, landmark-free co-registration of histology images and MSI data, respectively. MSI.EAGLE is freely available at github.com/Weljie-Lab-UPenn/MSI.EAGLE.

### Incorporation of spatial weighting during UMAP dimensionality reduction improves clustering

High-dimensional clustering of spatial data is heavily utilized as both hypothesis generating and hypothesis testing information by researchers. For MSI data specifically, Uniform Manifold Approximation Projection (UMAP)^21^ and Spatial Shrunken Centroids (SSC)^22^ are the two most commonly used methods. However, both methods begin to lose information when applied to MSI data at cellular and sub-cellular scales.

A series of standard UMAP analyses were applied to 5×5 µm, 2×2 µm, and 1×1 µm pixel size mouse brain samples acquired in positive ionization mode **(Fig. 3a)**. It is clear as we look Across this series, the nature of the spatial organization being captured changes fundamentally. At 5×5 µm, UMAP clusters correspond to gross anatomical regions, with little overlap in 2D space. At 2×2 µm, the clusters begin to become more diffuse, likely mapping to cellular or sub-cellular features. In the 1×1 µm data, the UMAP projection looks fragmented and uninterpretable with standard clustering methods, regardless of the UMAP parameters used for dimensionality reduction and clustering (**Fig 3a, b)**. However, it is clear from the localization of PC 38:4 and PE O-40:9 that gross structural information is contained within the 1×1 µm data, but UMAP analysis is unable to capture it **(Fig. 3c)**. Standard UMAP spectrally weights every pixel independently of its location in the tissue. At high enough spatial resolution, it is likely that cellular and sub-cellular features are now defining the resulting clusters, instead of regional metabolism. For example, the mitochondrial metabolism of each individual neuronal cell could be more similar to each other than the metabolism of a granule layer is to itself. When pixels are small enough to distinguish this mitochondrial metabolism, it now defines the UMAP cluster. The result of this is a UMAP that is unable to retain gross structural information. SSC, which weights spatial proximity of pixels, does the exact opposite. When applied to 1×1 µm data, SSC can clearly define sub-organ regions. However, it does not retain any of the cellular and sub-cellular information captured by UMAP within clusters **(Fig 3c).**

**Fig. 3.**
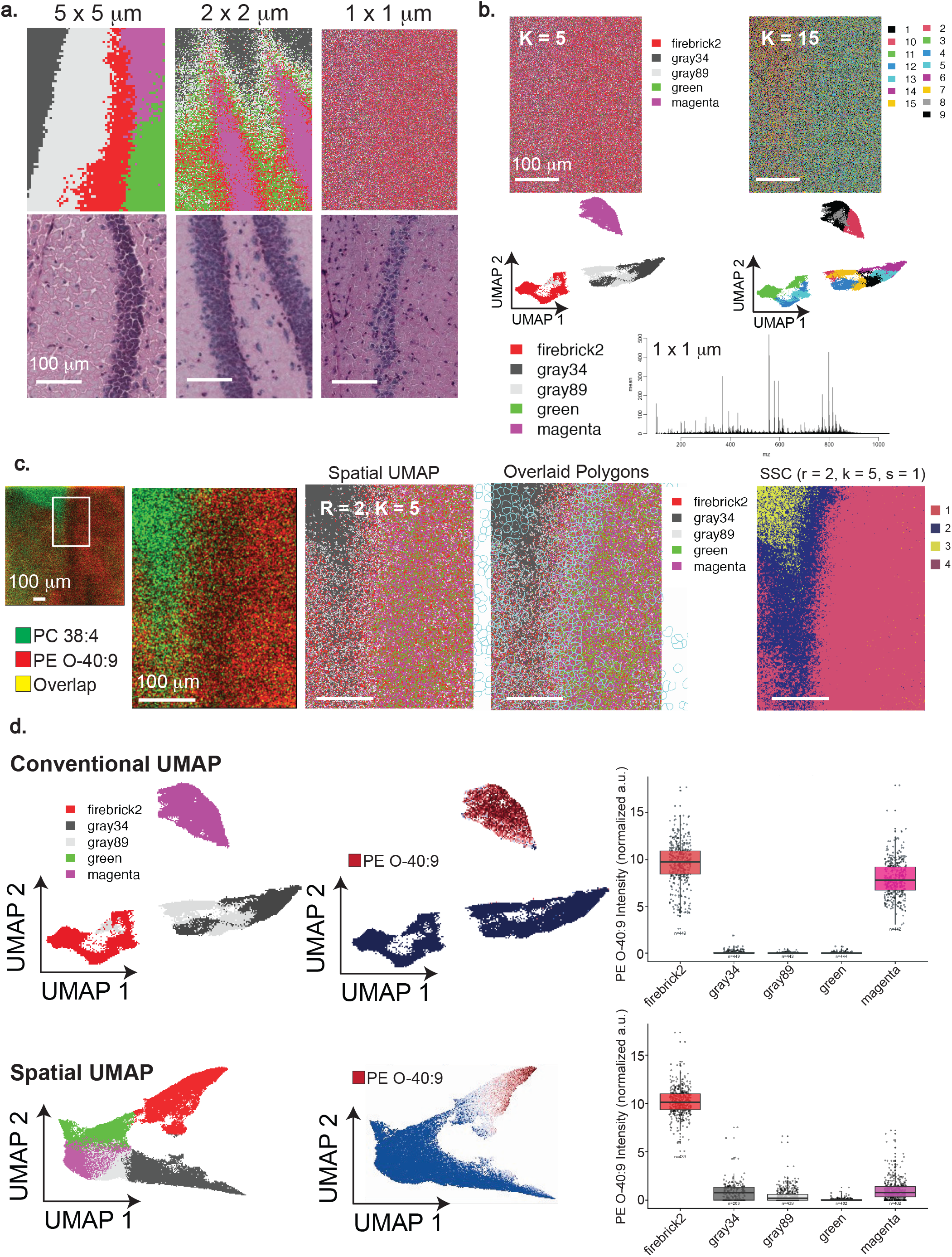
Spatially weighted UMAP preserves fine detail and gross structure at high resolution. Positive-mode DESI-MSI of the hippocampus/dentate gyrus region with post-acquisition H&E of the same section. **a,** Standard UMAP cluster maps (top) and matched H&E (bottom) at 5×5, 2×2 and 1×1 µm; clusters track anatomy at 5×5µm but disintegrate by 1×1 µm. **b,** Standard clustering of the 1×1 µm data at K = 5 and K = 15 with the corresponding UMAP embeddings and mean spectrum, showing that fragmentation persists across parameters. **c,** Ion images of PC 38:4 (green), PE O-40:9 (red) and their overlap (yellow) confirm intact tissue structure at 1×1 µm (overview with boxed ROI, and zoom), shown with the Spatial UMAP cluster map (R = 2, K = 5), the same clusters with overlaid cell-segmentation polygons, and SSC clusters with the same parameters (R = 2, K = 5). **d,** Conventional UMAP (top) versus Spatial UMAP (bottom): cluster embeddings, embeddings colored by PE O-40:9 intensity, and per-cluster PE O-40:9 box plots (median, IQR; whiskers 1.5x IQR; each point a pixel, n per cluster indicated). PE O-40:9 is confined to a single cluster (firebrick2) under Spatial UMAP but split between firebrick2 and magenta under conventional UMAP. Scale bars, 100 µm.

To this end, we developed an adaptation of the standard UMAP clustering algorithm. By integrating UMAP with Cardinal’s Spatial Fastmap, we produce an embedding that balances spectral similarity against spatial proximity. In a direct comparison between the conventional and spatial UMAP projections, we can see that in the spatial UMAP, spectrally similar pixels are still pulled together, but spatially contiguous regions are also encouraged to cluster together **(Fig 3c).** This prevents the embedding from fragmenting structures into disconnected clusters, while also retaining the fine detail present in high-resolution data. Additionally, spatial UMAP produces a more coherent metabolic profile per-cluster. Demonstrative of this is PE-O 40:9, which is elevated in only one cluster with spatial UMAP, but spread among two clusters with standard UMAP **(Fig 3d).** We quantify this gain in both spatial coherence and per-cluster coherence in **Supplementary Fig. 3**.

### Landmark-free Histology Fit enables cellular-level co-registration

Biological interpretation of subcellular MSI requires rigorous spatial registration with histological annotations and external molecular assays. MSI.EAGLE provides this registration framework within a module named Histology Fit (**Supplementary Fig. 2**). Unlike landmark-based co-registration strategies, this method quantitatively registers external images and/or annotations to MSI data in a common coordinate frame. In short, Histology Fit links histological signals derived from raw slide images to MSI ion intensity to determine the affine transformation necessary for accurate, quantitative co-registration **(Fig 4a).** A full description of the module details is available in the Supplementary Information.

**Fig. 4.**
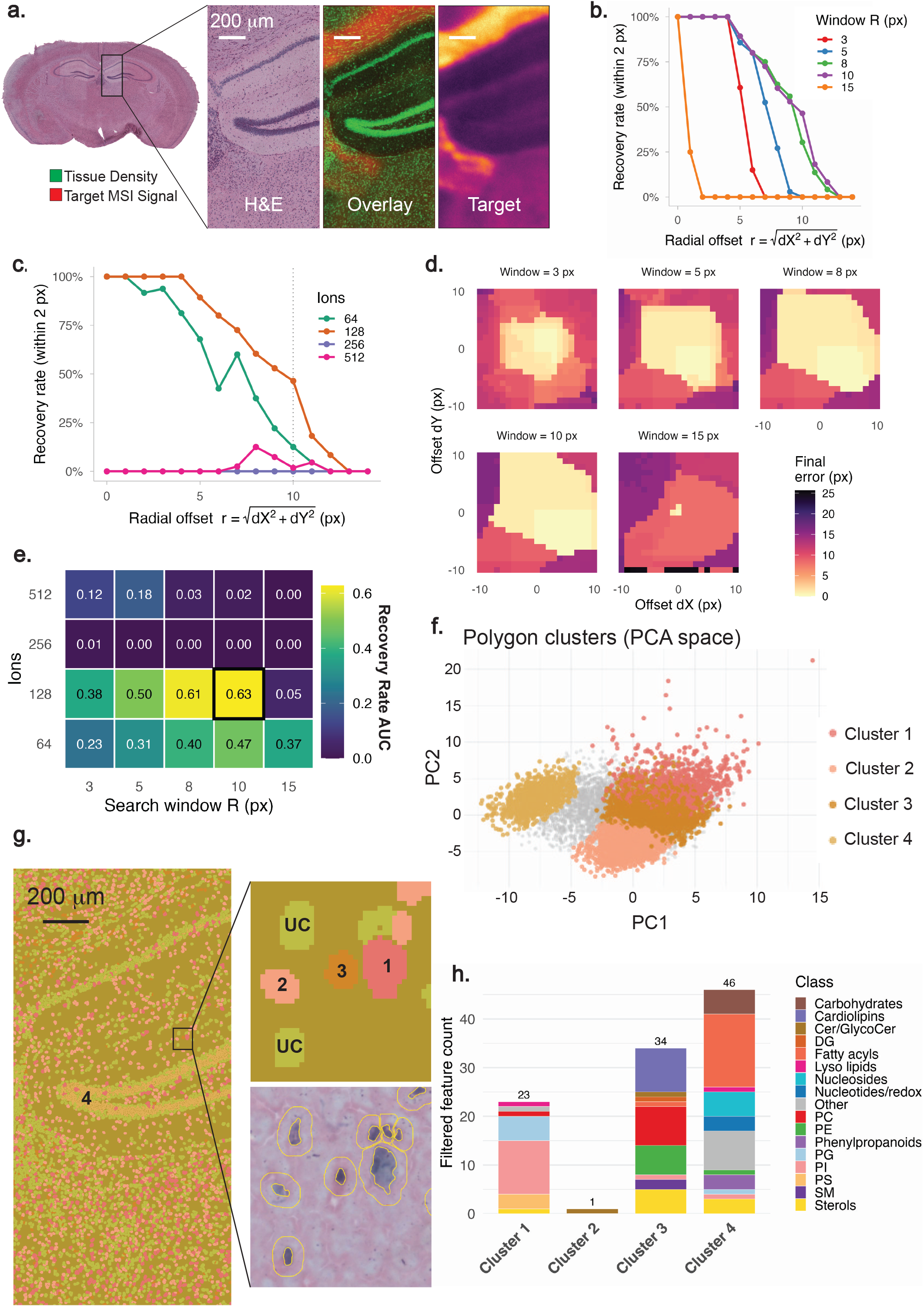
Landmark-free co-registration (Histology Fit) and morphology-resolved metabolomics. **a,** Histology Fit inputs for the section in Fig. 2b: H&E, the overlay of H&E-derived tissue density (green) and the MSI target signal (red), and the MSI target image; the affine transform aligning the two is solved from their correspondence. **b,c,** Registration recovery rate (return to within 2 px of ground truth) versus radial offset r = √(dX²+dY²) over a ±10-px, 441-point offset grid, colored by search-window radius R (**b**; 128 ions) and by number of informative ions (**c**; R = 10 px; dotted line, r = 10 px). **d,** Final registration error across the dX-dY offset grid for each search window (3-15 px). **e,** Recovery-rate AUC across ion count × search-window radius; the optimum (128 ions, R = 10 px; AUC 0.63) is boxed. **f,** Morphology-only clustering of segmented cells in PCA space (47 QuPath-derived features: nuclear diameter, staining intensities, nuclear-to-cytoplasmic ratio and 44 more), four clusters. **g,** Morphology clusters mapped onto the tissue with an enlarged inset (UC, unclassified) and the matched segmented H&E. **h,** Counts of differentiating metabolite features per morphology cluster, colored by compound class (Cluster 1, n = 23; Cluster 2, n = 1; Cluster 3, n = 34; Cluster 4, n = 46). Scale bars, 200 µm.

To place quantitative error bounds on the precision of the Histology Fit co-registration, we performed a synthetic offset-recovery experiment **(Fig 4b).** Known artificial translations were introduced to a converged reference registration of the same mouse brain sample from Fig. 2b, and the fit was re-run from each perturbed starting position. We then recorded whether and how accurately the registration returned to the true alignment as a function of user input parameters such as offset magnitude, search-window size and number of ions.

Encouragingly, the fit’s own converged optimum coincided exactly with the independently saved reference alignment (zero net displacement), confirming that the two agree. We then applied a ±10-pixel (±20 µm) grid of 441 artificial offsets and re-ran the fit from each. At the primary settings (128 informative ions, ±10-pixel search window), the registration recovered the reference alignment with high precision: displacements of up to 4 pixels (8 µm) were corrected in 100% of cases to a mean residual of less than one pixel (< 2 µm), and recovery remained above 50% out to 9 pixels (18 µm) before falling off as offsets approached and exceeded the search window **(Fig. 4b)**. Because 2×2 µm pixels place this recovery envelope at the single-pixel, single-cell scale - and because a user initializing the fit is realistically within a few pixels of the correct alignment - Histology Fit converges reliably to the correct registration in practice. Pooling recovery across all offset directions into a single recovery-versus-offset curve, we summarized precision as the area under this recovery-versus-magnitude curve (AUC); across the full grid of ion counts and search windows, the primary configuration achieved the highest AUC (0.63) **(Fig. 4e)**.

Recovery depended on the two principal user-set parameters. The number of ions used to build the MSI target signal showed a clear optimum at 128 informative ions (58% overall recovery, AUC 0.63); using fewer ions (64) modestly reduced recovery (AUC 0.47), whereas greatly increasing the ion count (256-512) diluted the structural PCA target with lower-variance, noisier signal and collapsed recovery almost entirely (AUC ≤ 0.02) **(Fig. 4c)**. The search-window size traded capture range against specificity **(Fig. 4d)**: windows that were too narrow (3-5 pixels) could not reach larger offsets, an 8-10-pixel window maximized the recovered region, and an overly wide window (15 pixels) admitted spurious off-tissue optima and degraded recovery sharply (AUC 0.05). Together these results define a robust operating point **(Fig. 4e)** - approximately 128 ions and an 8-10-pixel search window - and place a quantitative, direction-averaged precision bound of roughly one pixel (2 µm) on Histology Fit co-registration for offsets within the search window.

### Cell morphology in brain tissue is linked to metabolic state

With co-registered MSI and histology data, we performed morphological phenotyping of cells by their QuPath-derived physical attributes (nuclear diameter, staining intensities, nuclear-to-cytoplasmic ratio, and 44 additional parameters) using unsupervised PCA-based clustering, independent of any MSI data in MSI.EAGLE **(Fig 4f,g)**.^23^ Mapping these morphology clusters onto the MSI reveals that physically distinct cell populations have significantly different metabolic profiles. Specificity analysis, in which the normalized fold-change enrichment of each metabolite is calculated relative to the runner-up cluster, identified metabolite classes with strong cell-phenotype associations **(Fig 4h).**

Cluster 1 was made up of cells containing large cell bodies and prominent nuclei, consistent with neuronal morphology **(Fig 4g)**. Metabolically, the cluster was dominated by phosphatidylinositol, phosphatidylserine, and phosphatidylglycerol species **(Fig 4h).** Phosphatidylserine is highly enriched in neural membranes and supports membrane signaling, neurotransmission, and synaptic protein function, while phosphoinositides are major organizers of neuronal membrane identity and trafficking.^24,25^ A signaling-membrane-rich phenotype, coupled with their large morphology, indicate Cluster 1 may be a neuronal subpopulation, although the present data do not by themselves permit a definitive cell-type assignment.

While Cluster 2 does not appear to have a defining metabolic profile, Cluster 3 was enriched for a tightly related cardiolipin series (CL 72:5, CL 72:4, CL 72:3, CL 76:5, CL 72:2, CL 76:3, CL 68:2, CL 68:1, CL 78:3), together with phosphatidylcholine and phosphatidylethanolamine species. This pattern suggests that Cluster 3 corresponds to a mitochondria-rich cellular compartment.^14^ Cluster 3 was also enriched in several sulfatide lipids, a major lipid component of myelin and oligodendrocyte membranes.^26^ Both the cardiolipin and sulfatide enrichments indicate Cluster 3 contains a high proportion of oligodendrocytes/myelin.

Cluster 4 corresponds morphologically to the dentate gyrus **(Fig 4g)**. Among the four clusters, Cluster 4 carried the largest and most class-diverse panel of differentiating features (n = 46) and was distinguished by a pronounced fatty-acyl enrichment. Its highest-confidence features included N-arachidonoyl taurine, hydroxylated 20:4 and 22:6 fatty acids, an oxygenated DHA species (FA 22:6;O2), LPS 18:0 (lysophosphatidylserine), cytidine, and glycerophosphoglycerol, alongside xanthine and other small-molecule metabolites **(Fig 4h).** Taken together, this signature indicates active lipid turnover, remodeling, and mediator biology -- a pattern which fits the dentate gyrus’s role as a site of ongoing adult neurogenesis and high synaptic plasticity.^27^ Cytidine feeds the CDP-choline/CDP-ethanolamine (Kennedy) pathway for de novo phospholipid synthesis, while lysophosphatidylserine and glycerophosphoglycerol are phospholipid deacylation/remodeling intermediates; together these point to elevated membrane biogenesis and turnover, and xanthine is consistent with the high purine and energy turnover such a region would demand. We also see signatures indicative of bioactive-lipid signaling: N-acyl taurines are recognized endogenous FAAH-regulated lipid messengers, and the hydroxylated and oxygenated arachidonic-acid- and DHA-derived species are oxylipin-like mediators broadly implicated in inflammatory and vascular signaling in the nervous system.^28^ These mediator-type species are most plausibly contributed by the dentate gyrus’ non-neuronal compartment containing microglia, astrocytes, and vasculature, rather than by granule neurons themselves. This demonstrates the limitation of morphological cell definitions such as this; functionally and molecularly distinct cell populations can appear nearly identical with standard stains and/or analyses. Thus, the cell-type identifications presented here represent plausible, but still provisional, assignments.

### Cell-type-resolved metabolomics via predicted transcriptomic signal integration

To address this cell-type ambiguity, we integrated MSI.EAGLE with iSCALE,^29^ a machine-learning framework that predicts spatial gene expression (and by extension cell-type identity) directly from H&E images. iSCALE was trained on paired spatial-transcriptomics and histology datasets and produces cell-type probability maps at the resolution of the H&E image, which we verified against the Allen Brain Atlas^30–32^ **(Supplementary Fig. 4)**. MSI.EAGLE imports these iSCALE cluster maps and registers them to the MSI data using the common coordinate frame shared between the H&E image, the iSCALE clusters, and the MSI data.

We first registered the MSI data and H&E image using Histology Fit as described above, and verified the alignment at the sub-organ **(Fig. 5a)** and cellular **(Fig. 5b)** scales. MSI.EAGLE then imported and registered the externally generated iSCALE cell-type clusters using the same transformation parameters, yielding a quantitatively co-registered cell-type map suitable for downstream metabolic comparison **(Fig. 5c)**. To demonstrate cell-type-resolved metabolomics, we contrasted two spatially and transcriptomically distinct populations captured in the same section: ependymal/choroid-plexus astrocytes (astro-epen) lining the third ventricle, and glutamatergic neurons of the medial/lateral habenula (MH-LH glut).

**Fig. 5.**
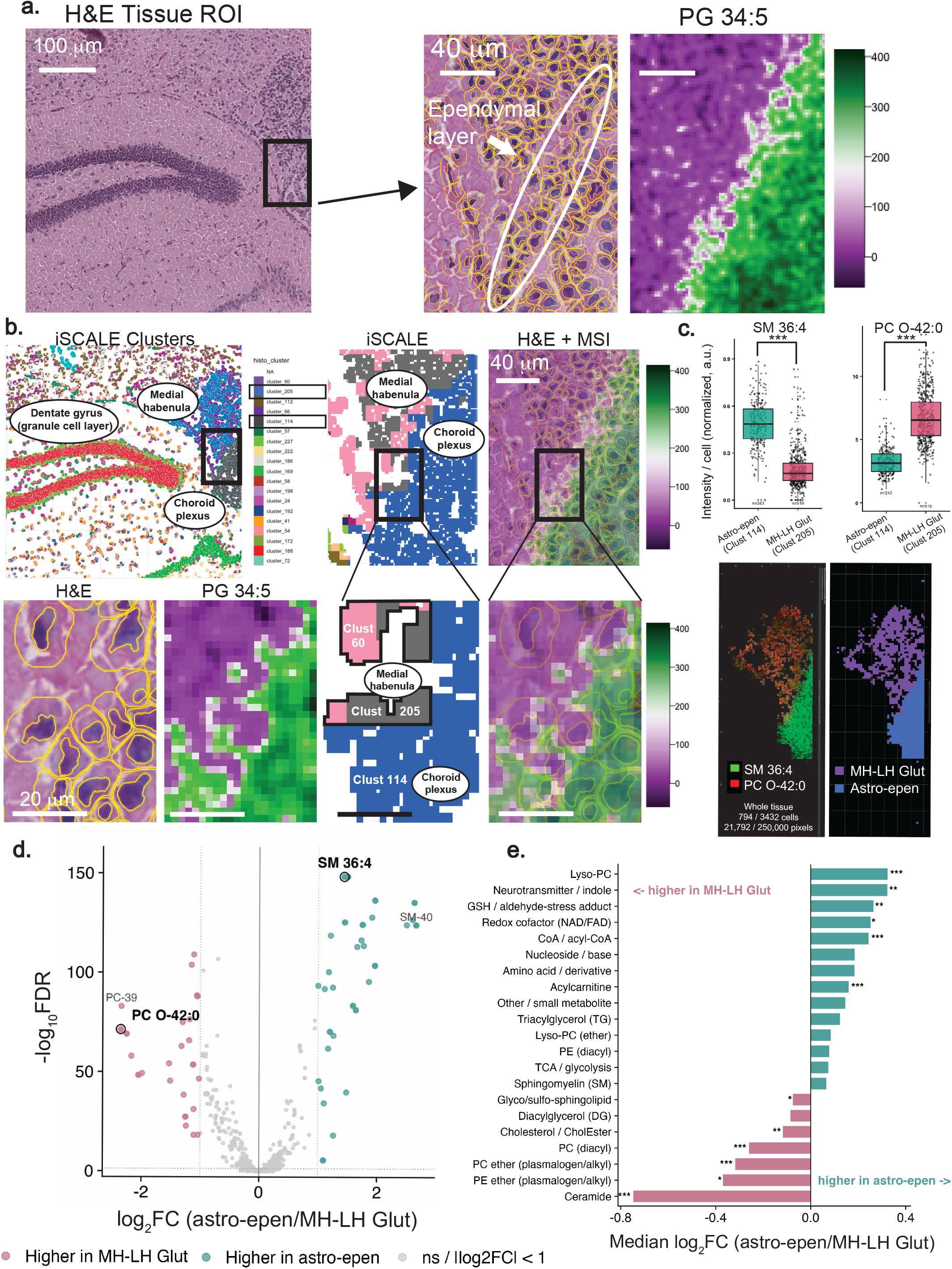
Cell-type-resolved metabolomics by integration of iSCALE-predicted transcriptomics. **a,** Cellular-scale verification of MSI-H&E registration: H&E ROI, a segmented zoom over the ependymal layer, and the co-registered PG 34:5 ion image. **b,** iSCALE-predicted histological clusters (dentate gyrus, medial habenula and choroid plexus annotated), the simplified iSCALE map and H&E+MSI overlay locating cluster 205 (medial-habenula glutamatergic neurons, MH-LH glut) and cluster 114 (choroid-plexus/ependymal astrocytes, astro-epen), enlarged H&E and PG 34:5 insets, and whole-tissue maps of SM 36:4 / PC O-42:0 and of the two assigned populations (794/3,432 cells; 21,792/250,000 pixels). **c,** Per-cell intensity of the marker features SM 36:4 (astro-epen) and PC O-42:0 (MH-LH glut) in the two populations (astro-epen, n = 243; MH-LH glut, n = 519 cells; box: median and IQR; ***). **d,** Volcano plot of the 829 co-registered targeted features: log₂ fold change (astro-epen / MH-LH glut) versus -log₁₀ FDR; 510 reach FDR < 0.05 and 80 exceed twofold (49 higher in astro-epen, teal; 31 higher in MH-LH glut, pink); dashed lines mark |log₂FC| = 1 and FDR = 0.05. **e,** Median log₂ fold change per compound class (≥4 annotated features); class enrichment assessed by two-sided Mann-Whitney U test of each class’s log₂FC against all other features (***P < 0.001, **P < 0.01, *P < 0.05). Scale bars: 100 µm (**a,** H&E ROI); 40 µm (**a, b** zooms); 20 µm (**b**, enlarged H&E).

Despite occupying adjacent regions of the same tissue, the two cell types were sharply separated by their metabolomes: of 829 co-registered targeted features, 510 differed at FDR < 0.05 and 80 exceeded a two-fold change (49 higher in astro-epen, 31 higher in MH-LH glut) **(Fig. 5d)**. Critically, the direction of these differences is consistent with the transcriptomically assigned identities rather than merely failing to contradict them.

The separation is dominated by membrane-lipid class **(Fig. 5e)**. MH-LH glut neurons are enriched in the canonical constituents of neuronal and synaptic membranes - ether-linked (plasmalogen) phosphatidylcholines and -ethanolamines (median log₂FC −0.32 and −0.37; MWU p = 1×10^−11^ and 0.03), polyunsaturated diacyl-PCs (−0.26; p = 1×10^−12^), and ceramides, the single most neuron-shifted class (−0.75; p = 4×10^−6^). Their strongest individual feature is the neuronal ether-lipid PC O-42:0 (log₂FC −2.35, FDR 5×10^−72^), followed by PC O-40 (−2.25, FDR 9×10^−70^), consistent with the known plasmalogen enrichment of neuronal membranes.^33^ Astro-epen cells instead carry the membrane signature of a secretory cerebrospinal-fluid (CSF) interface: lyso-phosphatidylcholines (median log₂FC +0.32; p = 6×10^−7^) and long-chain sphingomyelins, including the polyunsaturated SM 36:4 highlighted above (log₂FC +1.44, FDR 4×10^−149^) and SM 40:1/40:2 (+2.6, FDR < 10^−124^). Although the sphingomyelin class as a whole is heterogeneous, its large-magnitude members are strongly astro-epen-skewed (29 SMs significantly up in astro-epen vs. 4 in MH-LH glut) **(Fig. 5e)**.

Beyond structural lipids, astro-epen cells held higher pools of acyl-transfer and energy intermediates such as acyl-CoAs (median log₂FC +0.24; p = 5×10^−8^), acylcarnitines (+0.16; p = 1×10^−5^), and NAD/FAD redox cofactors (+0.25; p = 0.05), including acetyl-CoA (+0.36; FDR 2×10^−15^). These encompass the expected profile of a mitochondria-dense, high-throughput secretory epithelium engaged in CSF production **(Fig. 5e)**.^34^ They were also selectively enriched in glutathione-reactive-aldehyde conjugates (+0.26; p = 7×10^−3^), including the 4-hydroxynonenal-glutathione adduct (HNE-GSH; FDR 4×10^−4^), consistent with the choroid plexus’s role as the brain’s principal detoxification and blood-CSF barrier tissue **(Fig. 5e)**.^35^

Individual functional metabolites reinforced these class-level trends. The glutamate-glutamine pair behaved as a built-in positive control for the glutamate-glutamine cycle: glutamate itself did not differ between the two populations (log₂FC +0.08, ns), as expected for a ubiquitous metabolite, whereas glutamine (the product of glial glutamine synthetase) was significantly elevated in the glial astro-epen population (+0.27, FDR 0.01). This enrichment follows directly from the compartmentalization of the cycle: glutamine synthetase, the enzyme that amidates glutamate to glutamine, is expressed almost exclusively in astrocytes, so glutamate cleared from the synaptic space is converted to glutamine within the astrocytic compartment before glutamine is shuttled back to neurons to regenerate the neurotransmitter pool. Newly synthesized glutamine therefore accumulates preferentially on the astrocytic (astro-epen) side, whereas glutamate is held at comparable levels in both populations. A panel of CSF-borne neuro-metabolites (dopamine, melatonin, indole-3-acetic acid, nicotinamide) was likewise concentrated at the astro-epen/CSF interface (all log_2_FC +0.26 to +0.58, FDR ≤ 10^−3^ except IAA), the second-most astro-epen-shifted class overall **(Fig. 5e)**. Together, these data are consistent with our iSCALE cell-type assignments, as the metabolism differentiating these cell types independently regenerates their identities.

Also, this demonstration shows that our pipeline using only inexpensive MSI hardware modifications, a simple H&E stain, and open-source software, can resolve chemically coherent, cell-type-specific metabolic states from predicted-transcriptomic cluster maps.

## Discussion

Spatial metabolomics has lagged behind spatial proteomics and transcriptomics because both the acquisition and analysis of high-resolution data have remained expensive, bespoke, hardware-intensive, and code-dependent. Recent works have pushed mass spectrometry imaging to the cellular and subcellular scale, but each advance makes the tradeoff between resolution and these detracting factors explicit; Transmission-mode MALDI-2 combined with fluorescence microscopy built directly into the ion source has achieved genuinely single-cell and subcellular imaging, landmark-free co-registration of microscopy with MSI on the same section, and automated segmentation of tens of thousands of cells.^36^ This and other studies are compelling demonstrations of what subcellular spatial biology can deliver, but rely on highly specialized instruments and expertise that remain out of reach for most laboratories. Our aim here was to reproduce that same class of capability, with instrumentation and compute available to most labs, including: 1) subcellular sampling, 2) single-cell and subcellular metabolomics, 3) high-fidelity landmark-free co-registration, and 4) automated large-scale cell annotation. We demonstrate this can be done on stock commercial hardware with an inexpensive and reversible modification, and analyzed end-to-end in open-source, code-free software. In short, we sought to make subcellular spatial metabolomics cheap and easy enough to be broadly adopted.

The finest pixel sizes in the field belong to MALDI-2, which reaches ∼1 µm and below, but only by adding a secondary post-ionization laser and, in the most capable transmission-mode configurations, purpose-built source optics.^7^ Comparable resolution has also been approached from the ambient, DESI-adjacent direction, but again through specialized means: nano-DESI single-cell metabolomics relies on custom-fabricated tapered or microfluidic probes^37,38^ and expansion-based strategies such as 10x-DESI ExMSI reach an effective ∼5 µm only by embedding the section in a swellable gel and physically enlarging it, adding a substantial preparation step that departs from the native tissue state.^39^ Each of these is an impressive result, and each demands specialized probes, custom source geometries, or additional, sometimes tissue-altering, sample handling. We instead extended the low-flow DESI configuration of Towers et al. with two in-line C18 columns, additional fused-silica capillary, and a reduced emitter-to-sample distance - a change that is inexpensive, fully reversible, and installable on a stock DESI-XS source that already costs less than half of an equivalent MALDI platform.^12,13^ This was sufficient to reach 2×2 µm pixels routinely, and 1×1 µm in proof-of-principle, pixel sizes well matched to the 20-30 µm cells that make up most mammalian tissue, while preserving DESI’s native advantages: it is ambient, label-free, requires no matrix and no expansion, and leaves the tissue intact for downstream staining.

Also, these smaller pixels would be of little value if the spray smeared analyte beyond each sampled spot. At a tissue-glass boundary in 2×2 µm data, tissue-specific detection did not extend measurably onto the adjacent glass (Fig. 1 d-g, median one-sided spillover of 0 µm, with ±4.4 µm anchor uncertainty bounds). This indicates that the modified source samples at 2×2 µm without detectable lateral carry-over of tissue signal. The same experiment also shows why edge sharpness is a poor proxy for resolution in MSI: ions with different interface chemistry transition at different positions across one physical edge.

The biological payoff is not incremental; At 5×5µm, signal is averaged across each cell body and reports only cell-to-cell variation in total abundance; at 2×2 µm the same cardiolipin signal resolves into perinuclear puncta, and a systematic analysis across 6,073 cells recovered the expected cytoplasmic enrichment of mitochondrial cardiolipins, with polyunsaturated arachidonoyl species conversely trending nuclear. That this structure is compressed at 5 µm and emerges only at 2 µm confirms that the added resolution captures genuine subcellular organization rather than finer sampling of an averaged signal.

Co-registration of microscopy with MSI is the recognized bottleneck in this field, and it is where the cost of the current state of the art is most apparent. Aligning MSI with histology has traditionally meant either imaging adjacent sections (introducing well-documented registration error at the cellular scale) or staining after acquisition, where matrix removal and laser irradiation can damage the tissue, and compromise antibody binding. The elegant solution demonstrated recently is to move the microscope into the ion source, eliminating cross-modality fiducial markers and enabling same-section, high-fidelity registration; but this again requires specialized in-source optics that most groups cannot replicate.^36^ Dedicated registration software exists as an alternative (e.g., MSIr and general multimodal registration tools), yet these are single-purpose utilities separate from the analysis pipeline and often still rely on manually placed landmarks.^40^ Our approach reaches the same goal from two inexpensive directions. Because DESI is matrix-free and label-free, the identical section can be H&E stained after acquisition without matrix-removal damage, removing the need for adjacent-section registration entirely. Also, Histology Fit performs the alignment itself: a landmark-free, quantitative registration carried out purely in software, applicable to any MSI-microscopy pair and requiring no modification to the instrument. We bounded its precision empirically: a 441-point synthetic offset-recovery experiment defined a robust operating point (∼128 informative ions, an 8-10-pixel search window) that recovered the reference registration to roughly one pixel (2 µm), single-cell precision at our working resolution. The same “no fiducial markers, same section, high-fidelity, automated” standard set by dedicated in-source microscopy is thus met here with a routine stain and free software, and within the same environment used for the rest of the analysis.

Existing analysis options force a choice between capability and accessibility: SCiLS Lab (Bruker Daltonik GmbH, Bremen, Germany) is powerful but commercial and closed-source; Cardinal, on which we build, is open and rigorous but code-based and therefore inaccessible to non-programmers;^10^ and tools such as METASPACE^41^ (metabolite annotation) or M2aia^42^ (multimodal visualization and 3D reconstruction) each address one part of the workflow well, but must be stitched together for a complete analysis. MSI.EAGLE instead unifies import, clustering, co-registration, statistics, and figure export in a single vendor-agnostic, code-free interface, so that a dataset can be taken from raw .imzML format to publication-quality figures without writing a line of code or leaving the application. Within it, Spatial UMAP addresses a problem specific to high-resolution data: conventional UMAP fragments as pixels shrink, because a pixel small enough to isolate a subcellular compartment begins to group like compartments across the tissue rather than like anatomy within it. At 1×1 µm pixel size, the embedding becomes uninterpretable even though ion images confirm structure is present. By balancing spectral similarity against spatial proximity, our Spatial UMAP preserves both fine detail and gross structure and yields more coherent per-cluster profiles. Automated QuPath-based cell detection then supplies morphological annotations for thousands of cells at a time: the same automated, large-scale segmentation that specialized platforms achieve, obtained here with open tools on consumer hardware.

The clearest test of this accessible pipeline is whether it can resolve cell-type-specific metabolism, the capability that has, until now, depended on specialized instrumentation. The leading approaches pair MSI with a second, costly measurement: integration of imaging mass cytometry with MALDI-MSI assigns single-cell metabolic profiles to immunophenotyped cells, but requires a mass cytometry instrument and panels of metal-tagged antibodies^43^, while joint metabolite-protein frameworks such as scSpaMet add multiplexed antibody imaging to reach the same end.^44^ Both are elegant, but both make cell-type resolution contingent on antibody reagents and specialized hardware tuned to each study. Our pipeline reaches cell-type-resolved metabolism from a routine H&E stain and open-source iSCALE-predicted transcriptomics alone. This requires no antibodies and no second instrument on the imaged section. Critically, it does so while closing the loop from measurement back to biology.

Morphology-based clustering, computed independently of any MSI data, mapped physically distinct cell populations onto distinct metabolic profiles, while also exposing the limits of morphology alone: functionally distinct populations can be indistinguishable by standard stains, leaving such assignments provisional. Integrating iSCALE-predicted cell-type maps resolved this ambiguity, identifying ependymal/choroid-plexus astrocytes lining the third ventricle and habenular glutamatergic neurons. Additionally, the metabolome was consistent with these cell identities via neuronal plasmalogen and ceramide enrichment, a choroid-plexus signature of secretory, mitochondria-dense, detoxifying epithelium, and the glutamate/glutamine pair serving as a built-in positive control for the astrocytic glutamine-synthetase step. That an orthogonal modality reconstructs cell identities assigned from histology alone demonstrates that the entire chain (inexpensive hardware, a routine H&E stain, and open-source software) resolves real, cell-type-specific biology.

Several limitations remain, marking where more specialized methods retain genuine advantages. MALDI-2 and expansion-based imaging still surpass our method in outright spatial resolution, reaching the sub-micron and ∼500 nm regimes; our advantage is one of accessibility, label-free operation, and cost rather than a claim to the finest achievable pixel. While 1×1 µm data are acquirable, they remain difficult to cluster, and Spatial UMAP improves interpretability without fully solving unsupervised segmentation at that scale. Finer pixels also carry real acquisition costs, on the order of ten hours for a 1×1 mm section at 2×2 µm pixel size (Supplementary Table 1) and a smaller sampling area demands more instrument sensitivity. Thus, the practical resolution ceiling is set as much by throughput as by the source. Our spillover measurement also has defined bounds. It comes from a single section along one axis. It sets an upper bound on tissue-to-glass carry-over rather than measuring instrument resolution, and it cannot separate spread caused by cutting from spread caused by the DESI solvent. Distances are referenced to a molecular anchor because the optical tissue front sits about 8 µm from it, a difference in how the two edges are defined, and absolute optical registration is uncertain by about 9 µm.

Our cell-type assignments rest on predicted rather than directly measured molecular identity: unlike the antibody and mass-cytometry-based approaches above, which measure protein identity directly, iSCALE outputs are inferred from histology, and although concordant with the Allen Brain Atlas, they remain a hypothesis-generating layer that ultimately warrants validation against measured spatial transcriptomics. Likewise, our lipid- and metabolite-class assignments rest on mass accuracy without orthogonal structural confirmation, and Histology Fit assumes a well-behaved 2D affine relationship and a reasonable user initialization, so sections with substantial non-linear distortion fall outside the regime we characterized.

These limitations point directly to the next steps. Pairing this workflow with measured, rather than predicted, spatial transcriptomics or imaging-based single-cell methods would convert provisional cell-type assignments into confirmed ones and enable true multimodal, cell-resolved metabolic atlases. Because MSI.EAGLE consumes vendor-agnostic .imzML, the same analysis layer applies unchanged to MALDI, nano-DESI, and other platforms, so laboratories already invested in high-end instrumentation lose nothing by adopting it; the tool is meant to complement these specialized methods, not compete with them. Its modular, open-source design likewise invites community extension of clustering, registration, and statistical methods, and extending the approach across tissue types. Above all, by reducing subcellular acquisition to an inexpensive, reversible modification and its analysis to free, code-free software, this work places high-resolution spatial metabolomics within reach of any laboratory with a commercial MSI platform and a workstation, bringing its accessibility into line with the spatial proteomics and transcriptomics methods it complements.

## Experimental Section

### Animals and ethics

Mouse brains were obtained from wild-type mice, as described previously.^45^ Mice were given access to food and water ad libitum in filter-topped cages and given autoclaved food and water at the University of Pennsylvania University Laboratory Animal Resources (Penn ULAR) facility. All experiments were performed in accordance with the guidelines of Penn ULAR and were approved by the local institutional animal care and use committee (IACUC). At 10-14 weeks, animals were euthanized and organs were immediately harvested, flash-frozen in liquid nitrogen, and stored at −80 °C until sectioning.

### Cell culture

SNU-449 hepatocellular carcinoma cells (ATCC, CRL-2234, RRID:CVCL_0454) were maintained in DMEM supplemented with 10% fetal bovine serum and GlutaMAX. For imaging, cells were plated in 4-well cell-culture slides (CCS-4, MatTek) at 5×10⁵ cells per well and incubated in the same medium for 24 h to allow adherence to the slide surface. Medium was aspirated, wells were washed with phosphate-buffered saline (PBS) to remove residual medium, and slides were air-dried prior to MSI acquisition.

### Tissue-edge spillover analysis

Lateral carry-over of tissue signal onto glass was assessed at the edge of a mouse brain section imaged at 2×2 µm in negative ion mode (708 peak-picked features) The analyzed field spanned 201 image columns (x = 275-475; 402 µm along the edge) and 150 rows (300 µm) perpendicular to the edge. The primary readout was detection fraction, P(I > 0): the proportion of pixels in which an ion had nonzero intensity. Because it depends only on whether an ion is present, it does not depend on total-ion-current normalization. Detection fraction was computed for each row within 20 contiguous blocks of ten columns (the last block spanning eleven). Each ion’s profile was then normalized as F_det_ = (p - p_glass_)/(p_tissue_ -p_glass_), where p_glass_ and p_tissue_ are the mean detection fractions in far-glass (rows 183-203) and deep-tissue (rows 263-283) reference bands. Thus _Fdet_ = 0 marks the far-glass baseline rather than the absence of signal. Distance was referenced to a molecular anchor rather than the optical edge. The anchor was the row at which the detection fraction of a tissue-specific feature (feature 259) crossed 50% (row 233), and signed distance was d = 2 µm (row - 233), with negative values on glass. Sixty-seven tissue-positive ions were manually selected before the spillover statistic was computed and were reduced to one representative per isotope or related-feature cluster. They were grouped a priori as low mass (*m/z* 185-326, n = 19), higher mass (*m/z* 649-820, n = 26) and other masses (n = 22). One-sided spillover, S_20_, was defined for each block and ion as the distance onto glass of the 20% F_det_ crossing nearest the anchor. Crossings on the tissue side were scored as 0 µm. Group values are medians across block and ion S_20_ values, not crossings of the group-median curve. Within-section uncertainty was estimated from 2,000 hierarchical block-bootstrap replicates (seed 20260917). Each replicate resampled the same 20 blocks for every ion before taking the group median. Pointwise 95% intervals for the group curves (**Fig. 1g**) are the 2.5th and 97.5th percentiles of the bootstrap distribution. Anchor-placement sensitivity (±4.4 µm) was taken as the offset between the detection anchor and an independent intensity-based edge estimate (row 230.8), and is reported apart from the bootstrap intervals. The optical tissue front, estimated from the co-registered post-scan brightfield image, lay about 8 µm on the glass side of the anchor. This offset reflects the different definitions of the two edges, and absolute optical registration was underdetermined by about 9 µm. S_20_ is measured from the molecular anchor and does not depend on optical registration. S_20_ is an upper bound on tissue-to-glass carry-over, because spread from cutting and from the DESI solvent cannot be separated in this design.

### Tissue sectioning and slide preparation

Flash-frozen tissues were embedded in 12.5% (w/v) fish gelatin and cryosectioned at 10 µm on a Leica CM1950 cryostat (Leica Biosystems) with chamber and specimen temperatures of −20 °C; brain sections were cut coronally. Sections were thaw-mounted onto SuperFrost glass slides and stored at −80 °C until analysis. Immediately before acquisition, slides were brought to room temperature and dried under N₂ flow for 1 h.

### Low-flow DESI source modification

DESI imaging was performed on a Waters DESI-XS source coupled to a Xevo G2-XS QTOF mass spectrometer (Waters Corporation, Milford, MA, USA). To achieve subcellular pixel sizes, the low-flow, high-resolution DESI configuration of Towers et al.^9^ was extended. Whereas the original configuration replaced the standard PEEK solvent line with a 25-µm-i.d. fused-silica capillary routed through a single in-line C18 column to raise backpressure and confine the desorption plume, the present configuration incorporates two in-line C18 columns (2.1 × 150 mm, 1.6 µm) and 30 m of 25 µm i.d. fused-silica capillary tubing installed in series along the spray-solvent flow path. The emitter tip-to-sample distance was reduced relative to the stock geometry (1-2 mm) down to 0.25 mm. footprintCollectively, these additions further increased pump backpressure beyond a low-flow configuration, stabilizing the low-flow spray, while the shorter emitter-to-sample distance tightened the spray footprint (**Fig. 1a, Supplementary Fig. 1**). The modification requires no permanent alteration of the source, uses only commercially available consumables, and is fully reversible by restoring the original solvent line. No modification of the mass spectrometer was performed.

### DESI-MSI data acquisition

The mass spectrometer was operated in negative- and positive-ion modes in sensitivity mode (resolving power ∼20,000 FWHM) over *m/z* 50-1,200, with a scan time of 0.053s. In negative-ion mode the spray solvent was 98% methanol containing 50 pg µL^-1^ leucine enkephalin as a lock-mass reference ([M-H]⁻, *m/z* 554.2615); in positive-ion mode the spray solvent was 98% methanol containing 50 pg µL ^-1^ leucine enkephalin ([M+H]⁺, *m/z* 556.2771) and 0.01% formic acid. The spray-solvent flow rate was 250 nL min^-1^, and nebulizing N₂ gas was maintained at 600 L h^-1^. The sprayer was set at 75° to the sample surface. The heated transfer-line temperature was 50 °C for SNU-449 cell acquisitions, and 400 °C for mouse brain tissue. The capillary voltage was 0.6 kV in negative mode and 0.8 kV in positive mode.

Imaging regions and raster parameters were defined in High Definition Imaging (HDI) software (Waters). Pixel sizes ranged from 2×2 µm to 5 × 5 µm; 1×1 µm acquisitions were performed for proof-of-principle experiments. Unless otherwise stated, imaging runs used a 30 µm s^-1^ stage raster velocity. Acquisition parameters, feature counts, and run times for all datasets are summarized in **Supplementary Table 1**.

### Post-acquisition H&E staining

Because DESI is ambient, matrix-free and label-free, hematoxylin and eosin (H&E) staining was performed on the same section after MSI acquisition, eliminating the need for adjacent-section registration. Immediately after acquisition, sections were fixed in 10% neutral-buffered formalin for 10 min, rinsed in deionized water, stained with Mayer’s hematoxylin for 3 min, rinsed in running tap water, blued in Scott’s tap-water substitute for 1 min, and counterstained with eosin Y for 1 min. Sections were then dehydrated through graded ethanol (70%, 95%, 100%), cleared in xylene and coverslipped with a xylene-based mounting medium. Stained sections were imaged on a uScope MXII slide scanner (Microscopes International) with a 40x objective, giving a pixel size of 0.1138 µm.

### Data conversion and preprocessing

Raw acquisition data were centroided and lockmass corrected using Waters MassLynx AFAMM, then converted to .mzML and .imzML using MSConvert (ProteoWizard) and imzML Converter v1.3, respectively. Converted .imzML files were imported into MSI.EAGLE v0.1^11^ running under R v4.4.1. Peak picking was performed within MSI.EAGLE using the Cardinal peakPick() routine with the “qTof1” parameter preset for the Xevo G2-XS QTOF data. All data was peakpicked using a 25-ppm mass-accuracy window. Ion images were displayed with histogram-equalization contrast enhancement and Gaussian smoothing with identical display settings for the 5×5µm and 2×2 µm acquisitions compared in **Fig. 1b,c**. For the mouse-brain resolution comparison, feature detection used a signal-to-noise threshold of 20, yielding 2,086 features at 5×5µm and 4,696 features at 2×2 µm (Supplementary Table 1). All processing was performed on an M1 Ultra Mac Studio (128 GB RAM, 16 cores) using 20-100 processing chunks.

### Metabolite annotation

Features were annotated by accurate mass against LIPID MAPS^46^ and HMDB^47^ using a 25ppm mass tolerance window, and considering [M-H]^-^, [M+H]^+^, [M+Na]+, and [M-2H]^2-^ adducts. Lipid shorthand nomenclature follows the LIPID MAPS consensus. Annotations rest on accurate mass without orthogonal structural (MS/MS) confirmation and are reported at the corresponding annotation confidence level.

### Spatial UMAP segmentation

Unsupervised segmentation was performed in MSI.EAGLE. Standard UMAP^21^ embeddings were computed on per-pixel peak-intensity data using the uwot R package^48^. Embeddings shown in **Fig. 3a** used MSI.EAGLE’s default UMAP parameters: 8 nearest neighbors, minimum distance 0.02, cosine distance metric, no PCA pre-reduction, set_op_mix_ratio 0.15, search_k 1000, 100 trees, spread 1, local connectivity 1, repulsion strength 1.2, negative sample rate 10, learning rate 1, 600 epochs, and spectral initialization. Spatial UMAP embeddings (**Fig. 1b,c** and **Fig. 3c,d**) used the same underlying UMAP parameters, applied to a Cardinal spatial Fastmap projection with radius r = 2, 20 Fastmap components, Gaussian spatial weighting, and Euclidean distance metric, following the application defaults. Resulting embeddings were clustered using k-means. For comparison, spatial shrunken centroids (SSC)^22,49^ segmentation was performed with Cardinal’s spatialShrunkenCentroids function (parameters r =2, k=5, s=1, and adaptive weighting) (**Fig. 3c**).

### Histology co-registration (Histology Fit)

Co-registration of H&E microscopy and QuPath-derived annotations with MSI data was performed in the MSI.EAGLE Histology Fit module. Registration was modeled as a 2D affine transformation (translation, rotation, and scaling); the scaling factor was fixed to the physical ratio of microscopy to MSI pixel sizes (µm per pixel) so overlays were placed in real spatial units. An initial alignment was set manually using slider, numeric, and nudge controls with adjustable overlay transparency, then refined quantitatively. Quantitative refinement held scale and rotation fixed and searched a grid of candidate X/Y offsets (user-defined range and step) for the translation maximizing agreement between a histology-derived feature map (emphasizing hematoxylin-stained, chromatin-rich, tissue-dense regions) and an MSI molecular target. The MSI target could be a single ion, an RGB composite, the total ion current, a low-order principal component computed from the most spatially informative ions, or a pixel-metadata field. Each candidate offset was scored by a combination of rank correlation, normalized mutual information, and boundary agreement, evaluated in parallel across cores; score surfaces, contour maps, and overlays were reviewed before applying the best-scoring transformation. Two color-map-independent refinements were available: a statistical fit optimizing inside-versus-outside or cluster-group ion statistics (t-test or ANOVA), and an edge fit aligning polygon boundaries to the MSI tissue edge. Applying the transformation mapped each cell polygon (and, where a nucleus GeoJSON was supplied, the corresponding nuclear and cytoplasmic pixels) onto MSI coordinates and wrote the annotations into pixel metadata. Transformation parameters were saved for reproducibility and for transfer to externally generated cluster maps (see iSCALE integration).

### Registration precision (synthetic offset-recovery)

Co-registration precision was bounded empirically on the 2×2 µm mouse-brain section (Fig. 2b) with a headless port of the Histology Fit scoring and candidate-selection code that reproduces the application’s logic exactly. The MSI target was a principal-component image built from the N most informative ions (leading three PCs fused with weights based on variance and spatial coherence); the histology feature was H&E “darkness” (1 - luminance). Both images were Gaussian-smoothed (σ = 1 px), and the search step was 1 px. The reference optimum M was the fixed point of the fit started from the saved registration, which coincided with it (Translate X = 9, Y = 43 px). A ±10-pixel grid of 441 artificial X/Y translations (1-px steps) was applied around M, and the fit was re-run from each start for four ion counts (64, 128, 256, 512) and five search-window radii (3, 5, 8, 10, 15 px): 8,820 fits in total, with 128 ions and a 10-px window as the primary configuration. A start counted as recovered if the re-fit landed within 2 px of M. Recovery rate was computed per integer bin of radial offset magnitude r = √(dX² + dY²), and direction-averaged precision was summarized as the area under the recovery-versus-r curve (trapezoidal rule, normalized by the range of r so that AUC runs from 0 to 1). Per-axis RMS error was also recorded.

### Cell detection and morphological phenotyping

Cell detection was performed in QuPath v0.6.0 on the 40x H&E images. Cells were segmented with QuPath’s built-in nuclear-staining/size-based detection, manually verified, and adjusted where needed. Cell boundaries were exported as polygons in GeoJSON format together with per-cell morphological measurements (nuclear diameter, staining intensities, nuclear-to-cytoplasmic ratio, and additional parameters; 47 features total); where nuclear/cytoplasmic compartmentalization was required, nucleus boundaries were exported as a second GeoJSON. After Histology Fit registration, morphology-only unsupervised clustering was performed within MSI.EAGLE by k-means over a reduced principal-component space of the QuPath measurements, independent of any MSI data; cluster labels were transferred to pixel metadata.

### Perinuclear enrichment and specificity analyses

For subcellular localization (**Fig. 2c,d**), Nuclear and cytoplasmic pixel masks were defined from the registered nucleus and cell polygons of all 6,073 cells: nuclear pixels fell inside a nucleus polygon, and cytoplasmic pixels fell inside a cell polygon, but outside its nucleus. The cytoplasm-to-nucleus abundance ratio was computed for each of 73 high-confidence targeted compounds.

For morphology-linked metabolism (**Fig. 4h**), 1,304 negative-mode features were compared across the four morphology clusters with meansTest, with Benjamini-Hochberg correction. For each feature, the cluster with the highest mean (the “winner”) was compared with the second-highest (the “runner-up”). Specificity was quantified as (winner - runner-up)/winner, and an effect score as specificity × log₂(winner/runner-up) × -log₁₀ FDR. Features with uncertain annotations or isobaric conflicts within 0.002 Da were removed, and duplicate annotations were collapsed to the highest-scoring feature. Within each cluster, a feature was called differentiating if its effect score was at or above the 75^th^ percentile and its specificity and log₂(winner/runner-up) were each at or above the 60th percentile (up to 50 per cluster).

### Cell-type prediction and integration (iSCALE)

Cell-type identities were predicted from H&E images using iSCALE^29^ and the Allen Brain Atlas.^30^ iSCALE’s unsupervised clustering of H&E-derived predictions was applied directly. Each iSCALE cluster was then assigned a cell type manually by comparing its spatial distribution with the Allen Brain Atlas reference (**Supplementary Fig. 4**). MSI.EAGLE imported the iSCALE cluster maps and registered them to the MSI data using the transformation parameters obtained by Histology Fit for the same section (**Fig. 5a-c**). Group comparisons were performed in MSI.EAGLE with Cardinal’s meansTest. For cell-type-resolved metabolomics (**Fig. 5c-e**), 829 features from the laboratory’s curated, annotated metabolite list were compared between the astro-epen and MH-LH glut populations with the cell as the unit of analysis. P values were corrected with the Benjamini-Hochberg procedure (significance at FDR < 0.05); the asterisks in **Fig. 5c** show meansTest FDR. Class-level shifts were evaluated by Mann-Whitney U test on per-class log₂ fold-change distributions.

## Data and Code Availability

Installation instructions and source code for MSI.EAGLE are available for access here: https://github.com/Weljie-Lab-UPenn/MSI.EAGLE

All data in this work are publicly available for download here: 10.5281/zenodo.22943183

## AUTHOR INFORMATION

## Author Contributions

**DB** assisted in software development, acquired and analyzed all data included, and wrote the manuscript with input from all authors. **TOH** advised on analysis and manuscript preparation. **ML** and **MYL** generated iSCALE clusters. **GH** assisted in histology annotation. **OBP** assisted in software development. **FA, PK,** and **AS** advised on data collection and analysis. **GP** provided mouse brain samples. **TPG** provided SNU-449 cells. **GAF** advised on experiment design and analysis. **A.M.W.** analyzed data, developed the MSI.EAGLE software architecture, and conceived and supervised the project, providing funding. All authors reviewed and approved the final manuscript.

## Acknowledgement of Funding Sources

This work was supported by grants from the National Institutes of Health: 1R01NR018836 from the National Institute of Nursing Research (NINR), 1R01HL142981 from the National Heart, Lung, and Blood Institute (NHLBI), 5R01DK120757 from the National Institute of Diabetes and Digestive and Kidney Diseases (NIDDK), and P01CA165997 from the National Cancer Institutes (NCI).

## Supplementary Information

### MSI.EAGLE Modules

#### Data Import and Setup

The data import and setup tab provides users with functionality to import, visualize, and preprocess MSI datasets. MSI.EAGLE users can specify the directory containing raw imzML files and a working directory for result storage within the data import section. Files are selected through an interactive data table in the GUI. This step supports parallel processing, with users able to configure the parallelization mode and number of cores.

For peak picking, two predefined parameter sets are provided. “qTof1” is suitable for quadrupole time-of-flight instruments, while “HiRes” is designed for high-resolution instruments. Users have the flexibility to adjust these parameters based on their specific acquisition instrument and analysis type.

The data visualization module generates quick visualizations of the imported data, allowing users to verify acquisition and data conversion quality before proceeding to more in-masked analysis. A mass spectrum plot is created using a user-defined percentage of randomly sampled pixels, typically set to a low value. Users can interactively adjust the *m/z* and intensity ranges for this plot, helping them decide on relevant mass ranges for further analysis steps.

For testing purposes, the demo data section of this tab includes options to load pre-processed datasets from the CardinalWorkflows package. These datasets include samples such as human renal cell carcinoma and pig fetus cross-sections.

The data setup module is built on reactive programming principles in Shiny, ensuring efficient updates to outputs based on user inputs. This approach optimizes the application’s responsiveness by performing computationally intensive operations, like data import and peak picking of large datasets, only when necessary.

#### File Restore and Overview Analysis

The sidebar panel allows users to select from four main operations: 1) opening a previously peak-picked file, 2) peak picking raw files, 3) adding two imagesets with the same coordinates, or 4) adding two imagesets with the same peak list. These options enable users to choose the appropriate workflow for the stage and structure of their data.

For de novo peak picking, users can specify parameters such as the percentage of pixels to sample, signal-to-noise ratio (SNR), minimum peak frequency, and peak picking method (diff, sd, mad, quantile, filter, or cwt). The app utilizes Cardinal’s peakPick() function with these user-defined parameters to identify peaks in the MSI data.

The main panel displays a dynamic table of the processed runs, allowing users to select specific runs for further analysis. An interactive MSI plot is generated, which provides a visual representation of the peak-picked data. The module also includes functionality for saving the processed data as a .imzML directory and RDS file. This ensures that users can easily export their processed data for further analysis or future use, avoiding arduous re-processing steps.

#### Segmentation - Uniform Manifold Approximation and Projection (UMAP)

MSI.EAGLE implements unsupervised segmentation of mass spectrometry imaging data using Uniform Manifold Approximation and Projection (UMAP) dimensionality reduction.^48^ Users can remove background from the pixel data, annotate anatomical regions through an integrated pData/UMAP editor, or clean up stray pixels in a dedicated mode. The UMAP algorithm is applied to the selected dataset using the uwot R package.^21^ Key parameters, including the number of nearest neighbors, minimum distance, distance metric, number of trees, and an optional PCA pre-reduction, can be adjusted through the GUI.

UMAP analysis is performed on the pixel intensity data, and the resulting low-dimensional UMAP embeddings are then clustered using one of ten algorithms. K-means is the default method, but options include: hierarchical, DBSCAN, HDBSCAN, Spectral Clustering, K-medoids, Fuzzy C-means, Model-based Clustering (Mclust), Self-Organizing Map (SOM), and Spherical K-means. Users can interactively visualize the UMAP results and clustering, with options to display reduced color representations or cluster assignments. The GUI allows selection of specific clusters or colors to refine the segmentation.

Processed segmentation results can be stored and applied to the full dataset. For anatomical segmentation, users can annotate specific regions by selecting clusters and assigning labels, and a dedicated mode removes isolated pixels to clean up the segmentation. Processed and annotated data can be saved for downstream analysis as both an .imzML directory and .rds file in the working directory, and long-running operations can be autosaved to guard against data loss.

The segmentation tab leverages the Cardinal package for handling and visualizing mass spectrometry imaging data structures. Interactive plots are generated using ggplot2 and rendered as PNG images for presentation and/or publication by the user. This segmentation approach enables flexible, data-driven partitioning of imaging mass spectrometry data without relying on predefined anatomical boundaries. The interactive interface also allows iterative refinement of segmentation results, introducing tolerance for sub-optimal parameter selection by the user.

#### Segmentation - Spatial Shrunken Centroids (SSC)

MSI.EAGLE also utilizes the spatialShrunkenCentroids function from the Cardinal package to perform unsupervised segmentation of MSI data. This approach, based on the shrunken centroid method, incorporates spatial information to identify regions with similar molecular profiles.

The GUI allows users to select preprocessed datasets for analysis and set key parameters for the spatialShrunkenCentroids algorithm, including the spatial neighborhood radius (r), number of clusters (k), shrinkage factor (s), and the spatial weighting scheme (adaptive or Gaussian). Multiple parameter combinations can be evaluated in parallel. The segmentation results are cached to avoid redundant computations. Again, there is an optional step to remove isolated pixels is available through the fix_pix function to clean the image data.

Segmentation results are visualized using Cardinal’s image and plot functions. The GUI provides options to select specific segmentation models and color schemes for visualization. Users can interactively explore different cluster assignments and evaluate corresponding weighted spectra and statistics. The module allows storing and exporting the processed data, including segmentation results, for further analysis or reporting. All processing steps maintain the spatial and spectral integrity of the imaging data structure as implemented in Cardinal.

This segmentation approach enables users to objectively differentiate background ionization from sample data and identify regions of interest in mass spectrometry imaging experiments, facilitating the discovery of spatially-resolved molecular patterns in complex biological samples.

#### UMAP Embedding

MSI.EAGLE’s UMAP Embedding module enables interactive visualization and exploration of mass spectrometry imaging data in reduced dimensional space. Users select *m/z* ions of interest and color variables from pixel metadata through dropdown menus, and the module renders the UMAP projection for real-time interaction. The embedding visualization supports both intensity-based coloring of selected *m/z* values and categorical coloring based on metadata variables, with the color palette selectable from a broad set of sequential and diverging options.

For intensity-based visualization, the module implements a color scaling system with an adjustable midpoint value and optional log transformation. Points in the UMAP space can be selectively masked based on metadata values, with unselected points appearing in a neutral gray, and embedding colors can optionally be matched to those of the original UMAP segmentation. Point size and other visual parameters can be adjusted through interactive controls. A complementary spatial plot shows the selected *m/z* value or metadata variable mapped back onto the original image coordinates. All visualization options can be exported as high-resolution images for presentation or publication.

The module also supports re-clustering directly on an existing UMAP embedding. Any of the ten clustering algorithms available in the segmentation module can be applied to the embedding coordinates, with adjustable numbers of clusters, minimum points, and DBSCAN neighborhood radius. Memory-intensive methods are protected by configurable safety guards, which can be overridden and logged as diagnostics when needed. Cluster assignments are written to new pixel-metadata columns, and the resulting embedding clustering can be stored in the processed data and exported as an .imzML file for downstream analysis.

#### Phenotyping

The phenotyping tab of MSI.EAGLE enables users to associate phenotypic data with MSI datasets. Users can upload a tab-delimited text file containing phenotype information via a file input interface. The uploaded file is read using the read_samples function, which automatically detects the file format. Users can choose between two types of MSI images for phenotyping: 1) images restored from a.imzml or .rds file or 2) stored data from the application memory.

The application offers four phenotyping methods: 1) Spectral density, 2) Periodicity, 3) Breaks between samples, and 4) Manual (x & y limits specified in file). For the “breaks” method, users can specify a threshold value. The pixDatFill_mult function is used to perform the phenotyping for the first three methods, while pixDatFill_manual is used for the manual method.

After phenotyping, the results are displayed in an interactive table using the DT package. Users can select which columns from the phenotype data to include in the pData object. The application also provides an option to create interaction terms between two selected phenotype variables. Users can also partition the image into rectangular spatial tiles, specified by either tile size or tile count, and store the resulting tile assignments as a pData variable for downstream grouped analysis. The phenotyped data is visualized using Cardinal’s image function, allowing users to select which phenotype variable to display. The resulting image is rendered as a PNG file and displayed in the user interface. Users can save the phenotyped data, which includes the MSI image data with updated pData and fData objects.

This phenotyping module provides a user-friendly interface for associating phenotypic data with MSI datasets, enabling researchers to explore relationships between molecular profiles and sample characteristics within the application.

#### Masked Analysis

The Masked Analysis tab allows users to perform more comprehensive peak picking and extraction of lower abundance features. Users first select a previously segmented dataset file as a template for coordinate extraction. Coordinate extraction is performed by iterating over the run names in the segmented file and extracting corresponding coordinates using Cardinal’s coord function. The module then offers three analysis modes: untargeted peak picking, targeted peak binning, and mean spectrum-based peak picking.

For untargeted analysis, peak picking is performed using a custom HTS_reproc function, which incorporates multiple Cardinal functions. Targeted analysis utilizes Cardinal’s peakBin function, allowing users to input a list of annotated exact masses for binning. The mean spectrum approach combines Cardinal’s summarizeFeatures, normalize, peakPick, peakAlign, and peakFilter functions to calculate an average spectrum, perform peak picking, and use the resulting peak list to bin the raw data. Parallel processing capabilities are incorporated to handle large datasets, utilizing the bplapply function for certain operations.

After processing, the resulting peak-picked or binned data is visualized using Cardinal’s image function. The processed data can be saved for further analysis. This functionality enables researchers to extract and analyze depth-specific molecular information from complex samples in an interactive and user-friendly manner.

#### Statistics

The Statistics tab enables users to perform various statistical tests on the processed MSI data. The GUI allows selection of the analysis type, data grouping, and visualization options. Statistical methods implemented include means tests, spatial shrunken centroids (SSC), and spatially-aware Dirichlet Gaussian mixture models (spatial DGMM).

For means tests, the app utilizes Cardinal’s meansTest function to compare ion intensities between user-defined groups. Results are displayed as an interactive table with adjustable false discovery rate (FDR) thresholds. The spatial method (SSC) incorporates pixel coordinates to identify spatially-localized differences between groups. A spatially-aware Dirichlet Gaussian mixture model (spatial DGMM) is also available, modeling each group’s intensity distribution as a mixture of spatial Gaussian components.

MSI.EAGLE generates customizable visualizations of statistical results, including boxplots, means plots, and ion images colored by group membership or statistical significance. Users can interactively explore results, adjust plot parameters, and export tables and figures. Significant features can also be exported as ready-to-use input files for pathway and enrichment analysis in Mummichog and MetaboAnalyst.

The modular design and open-source nature of MSI.EAGLE allows easy addition of new statistical tests and visualization options. All analyses are performed server-side using R, with results passed to the user interface for display and interaction. This architecture enables efficient analysis of large imaging datasets within memory constraints.

#### Heatmap

MSI.EAGLE provides interactive heatmap visualization through a dedicated tab. Users can select variables for heatmap generation from phenotypic data associated with the MSI dataset. The heatmap is generated using the pheatmap R package, with options for row/column scaling, clustering methods, distance metrics, and color schemes.

Significance filtering can be applied to show only features meeting a user-defined threshold based on statistical results. Row and column annotations can be added to provide additional context. The resulting heatmap can be customized through various parameters including clustering cutoffs, label display, and color palettes.

All visualizations can be exported as high-resolution PDF files. Associated data matrices (raw and scaled) are saved as tab-separated text files to facilitate further analysis. This functionality enables iterative exploration and quality control of high-dimensional mass spectrometry imaging data within an interactive framework.

#### Colocalization Analysis

In the Colocalization Analysis tab of MSI.EAGLE, options include selecting correlation variables and *m/z* values, numeric inputs for specifying the number of colocalized features and top correlations to plot, and options for choosing the ranking metric (Pearson correlation, Manders overlap coefficient, Manders colocalization coefficients M1 and M2, or Dice similarity).

Upon triggering the analysis, the colocalization analysis is performed using the colocalized function from Cardinal. This analysis considers the user-selected *m/z* value, number of features, and sorting method. Visualization of the colocalization results is achieved using Cardinal’s image function. The resulting plot displays the top correlated features as specified by the user, with options for contrast enhancement, normalization, and color scale customization. This implementation allows for interactive exploration of colocalization patterns in mass spectrometry imaging data, providing users with flexible options for analysis and visualization.

#### Co-Registration of H&E microscopy images with MSI data

Cell detection is performed outside of the MSI.EAGLE application, while co-registration of the resulting annotations with the MSI data is performed within it. H&E images are acquired using a uSCOPE MXII microscope at 40X and imported to QuPath for further processing. Using QuPath’s built-in cell detection algorithms, cells are automatically identified based on nuclear staining intensity and cell size parameters, manually verified, and adjusted where needed. Cell region boundaries are exported as polygons from QuPath and stored as GeoJSON files, together with their per-cell morphological measurements; where a distinction between nuclear and cytoplasmic compartments is required, nucleus boundaries are exported as a second GeoJSON file.

MSI.EAGLE co-registers the histology image and its QuPath annotations with the MSI data in a dedicated module, Histology Fit. Users load the H&E image, the polygon GeoJSON file (and, optionally, a matching cluster-color image or nucleus GeoJSON), and the processed MSI data (.imzML or .rds format), which may be the dataset already held in application memory or a separately uploaded file. The MSI target can be displayed as a single *m/z* ion, an RGB composite of two to three ions, the total ion current, or a pixel-metadata field, with adjustable intensity transformation and color palette. Registration is defined by an affine transformation combining translation, rotation, and scaling, where the scaling factor can be fixed to the physical ratio of the microscopy and MSI pixel sizes (µm per pixel) so that the overlay is placed in real spatial units. Users set an initial alignment with slider, numeric, and nudge controls while adjusting the overlay transparency to visually verify alignment, and then refine the transformation quantitatively.

The Histology Fit routine performs this quantitative refinement by holding the current scale and rotation fixed and searching a grid of candidate X and Y offsets, over a user-defined range and step, for the translation that maximizes agreement between a histology-derived feature map and an MSI molecular signal. The histology feature map emphasizes hematoxylin-stained, chromatin-rich, or tissue-dense regions, and the MSI target can be the current display, a fused image built from the most spatially informative ions, a low-order principal component computed from those ions, or a pixel-metadata field. Each candidate offset is scored by a combination of rank correlation, normalized mutual information, and boundary agreement between the two modalities, with the grid evaluated in parallel across cores. The resulting score surface, contour map, and target overlay are displayed as diagnostics, allowing candidate registrations to be reviewed before the best-scoring transformation is applied. Two complementary, color-map-independent refinements are also available: a statistical fit that optimizes the offset using inside-versus-outside or cluster-group ion statistics (t-test or ANOVA), and an edge fit that aligns polygon boundaries to the MSI tissue edge.

Applying the transformation maps each polygon (and, when a nucleus GeoJSON is supplied, the corresponding nuclear and cytoplasmic pixels) onto the MSI coordinates and writes the cell annotations into the pixel metadata. Polygons can optionally be grouped before mapping by clustering their QuPath morphological measurements, or geometry-only features for manually drawn polygons, using k-means over a reduced principal-component space, with the resulting cluster labels transferred to the pixel metadata. The co-registered data is exported as both .rds and .imzML files compatible with MSI.EAGLE, with the cell annotations preserved in the pixel metadata. Two tables are also exported: a region measurements table containing morphological features extracted from QuPath for each cell (including area, perimeter, and circularity), and a transformation parameters table recording the specific scaling, rotation, and translation values used for registration; these parameters can also be saved and reloaded to reproduce or transfer a registration.

### Quantitative Benchmarking of Spatial UMAP

The reduction in fragmentation and gain in per-cluster metabolic coherence attributed to spatial UMAP in **Fig. 3c,d** were illustrated qualitatively, using representative ion images and a single lipid species (PE O-40:9). To quantify these effects across the full 1×1 µm hippocampal dataset, we benchmarked conventional UMAP, spatial UMAP (R = 2), and SSC (R = 2) (each clustered at K = 5, matching **Fig. 3c,d**) against an independent histological reference: the 449 H&E cell polygons detected in QuPath and registered to the MSI pixel grid via Histology Fit (**Supplementary Fig. 3a**). Coloring each embedding by whether a pixel falls inside or outside a registered polygon confirmed that neither method resolves the cell compartment as a discrete region of embedding space (**Supplementary Fig. 3b**), so we scored agreement at the level of individual cells instead.

Because this reference marks where each cell is but not its type, we scored two cell-resolution statistics, each corrected against a label-permutation chance level: within-cell purity (the fraction of a cell’s pixels sharing its single most common cluster label) and the effective number of distinct cluster labels per cell, an inverse-Simpson count that, unlike a raw tally of distinct labels, does not saturate once a cell spans dozens of pixels. Spatial UMAP increased chance-adjusted within-cell purity relative to conventional UMAP (0.214 vs. 0.010) and reduced the effective labels per cell from 4.36 to 2.95, generalizing across the full dataset the single-lipid fragmentation example in **Fig. 3d** (**Supplementary Fig. 3c**). Spatial coherence, measured by Moran’s I of the cluster labels, rose from 0.004 to 0.271 over the same comparison, and the fraction of adjacent pixel pairs sharing a cluster label from 0.219 to 0.435 (**Supplementary Fig. 3c**). This ranking held at every clustering granularity tested from K = 3 to K = 10 (**Supplementary Fig. 3d**).

SSC scored highest of the three methods on every cell-level and spatial-coherence metric (purity 0.635; Moran’s I 0.821), but at the cost of spectral discriminability: its spectral silhouette (cosine distance, in the original peak-picked feature space) was the lowest of the three (−0.054), consistent with the coarse regional partition it produces at this resolution (**Supplementary Fig. 3a**) and with the loss of cellular and sub-cellular detail described for SSC in **Fig. 3c**. Notably, the H&E cell segmentation itself has a silhouette near zero (−0.0004), showing that true cell boundaries are not spectrally separable at the pixel level either; the higher silhouette of conventional UMAP (+0.105) therefore reflects spectrally compact but histologically uninformative clusters rather than a genuine advantage over spatial UMAP (**Supplementary Fig. 3c**).

We additionally computed the adjusted Rand index of each method’s clustering against a whole-field cell/non-cell mask; this was at chance (|ARI| < 0.01) for every method, because the non-cell compartment is dense neuropil rather than empty space and is not recoverable from a five-way partition of pixel intensities alone. Parameter sensitivity was assessed by repeating the analysis over a grid of spatial radii (R = 0-4) and FastMap component counts (2-20) (**Supplementary Fig. 3e**). Both cell-level purity and spatial coherence increased with R across this range without turning over (best-in-row chance-adjusted purity 0.067 at R = 0 rising to 0.441 at R = 4), while the spectral silhouette fell from 0.101 to 0.016 over the same span; performance was insensitive to component count above two (0.237-0.256 across 3-20 components at R = 2, versus 0.179 at two components). Plotting the two axes against one another places every parameter setting on a single monotonic front, with R = 2 mid-front — a conservative choice rather than one tuned to maximize either metric (**Supplementary Fig. 3f**). Substituting purely spatial (gaussian) for spectrally adaptive neighbourhood weights advanced further along that same front, raising purity and coherence at R = 2 to 0.432 and 0.675 but roughly quintupling the spectral-silhouette cost (−0.042 vs. −0.009) (**Supplementary Fig. 3f**).

---

**Supplementary Table 1:** Mouse Tissue Data and Analysis Characteristics.

| Data Characteristic | SNU-449<br>(5x5 µm) | SNU-449<br>(2x2 µm) | Spillover Brain<br>(2x2 µm) | Brain<br>(2x2 µm) | Brain<br>(5x5 µm) | Brain<br>(1 x 1 µm) | iSCALE Brain<br>(2x2 µm) |
| --- | --- | --- | --- | --- | --- | --- | --- |
| Pixel area | 25 µm<sup>2</sup> | 4 µm<sup>2</sup> | 4 µm<sup>2</sup> | 4 µm<sup>2</sup> | 25 µm<sup>2</sup> | 1 µm<sup>2</sup> | 4 µm<sup>2</sup> |
| Pixel dimensions<br>(h x w) | 5x5 µm | 2x2 µm | 2x2 µm | 2x2 µm | 5x5 µm | 1 x 1 µm | 2x2 µm |
| Total area acquired | 4,000,000 µm<sup>2</sup> | 4,000,000 µm<sup>2</sup> | 1,000,000 µm<sup>2</sup> | 2,000,000 µm<sup>2</sup> | 2,990,000 µm<sup>2</sup> | 1,000,000 µm<sup>2</sup> | 1,000,000 µm<sup>2</sup> |
| Number of pixels | 160,000 | 1,000,000 | 250,000 | 500,000 | 119,600 | 1,000,000 | 250,000 |
| Raster rate | 20 µm/second | 20 µm/second | 10 µm/second | 30 µm/second | 30 µm/second | 30 µm/second | 30 µm/second |
| Scan time | 0.236 seconds | 0.086 seconds | 0.186 seconds | 0.053 second | 0.153 second | 0.019 seconds | 0.053 second |
| Acquisition Time | 733 minutes | 1833 minutes | 916 minutes | 646 minutes | 369 minutes | 716 minutes | 362 minutes |
| Ionization Mode | Positive | Positive | Negative | Negative | Negative | Positive | Positive |
| Raw .imzML file size | 0.52 Gb | 3.20 Gb | 0.83 Gb | 1.71 Gb | 408 Mb | 3.41 Gb | 0.84 Gb |
| Raw .ibd file size | 13.7 Gb | 28.8 Gb | 16.93 Gb | 9.55 Gb | 6.60 Gb | 16.74 Gb | 13.56 Gb |
| Number of features<br>detected after untargeted<br>peak-picking (on-sample,<br>30 ppm window, 20 S/N) | 4,176 | 5,167 | 9,315 | 4,696 | 2,086 | 5,923 | 2,840 |
| Estimated mass<br>resolution | 7.5 ppm | 11.5 ppm | 9.5 ppm | 12 ppm | 9.5 ppm | 14 ppm | 9 ppm |

| Processing Times<br>(all pixels) | SNU-449<br>(5x5 µm) | SNU-449<br>(2x2 µm) | Spillover Brain<br>(2x2 µm) | Brain<br>(2x2 µm) | Brain<br>(5x5 µm) | Brain<br>(1 x 1 µm) | iSCALE Brain<br>(2x2 µm) |
| --- | --- | --- | --- | --- | --- | --- | --- |
| Initial peak-pick (full<br>area, 100 *m/z* range) | 4 minutes, 53 seconds | 19 minutes, 6 seconds | 3 minutes, 15 seconds | 2 minutes, 56 seconds | 55 seconds | 6 minutes, 21 seconds | 2 minutes, 52 seconds |
| Untargeted peak-picking<br>(full *m/z* range) | 22 minutes, 2 seconds | 156 minutes, 21 seconds | 35 minutes, 32 seconds | 49 minutes, 31 seconds | 5 minutes, 11 seconds | 113 minutes, 53 seconds | 43 minutes, 26 seconds |
| Targeted peak-picking<br>(full *m/z* range, ~200<br>targets) | 13 minutes, 47 seconds | 47 minutes, 39 seconds | 18 minutes, 13 seconds | 17 minute, 50 seconds | 4 minutes, 12 seconds | 40 minutes, 25 seconds | 16 minutes, 14 seconds |
| QuPath Cell Detection<br>and Polygon Export | 1-2 minutes | 1-2 minutes | 1-2 minutes | 1-2 minutes | 1-2 minutes | 1-2 minutes | 1-2 minutes |
| MSI + polygon<br>registration | 15-20 minutes | 15-20 minutes | 15-20 minutes | 15-20 minutes | 15-20 minutes | 15-20 minutes | 15-20 minutes |

**\*All processing done on an M1 Ultra Mac Studio with 128 Gb RAM, 16 cores, and between 20-100 chunks**

**Supplementary Figure 1.**
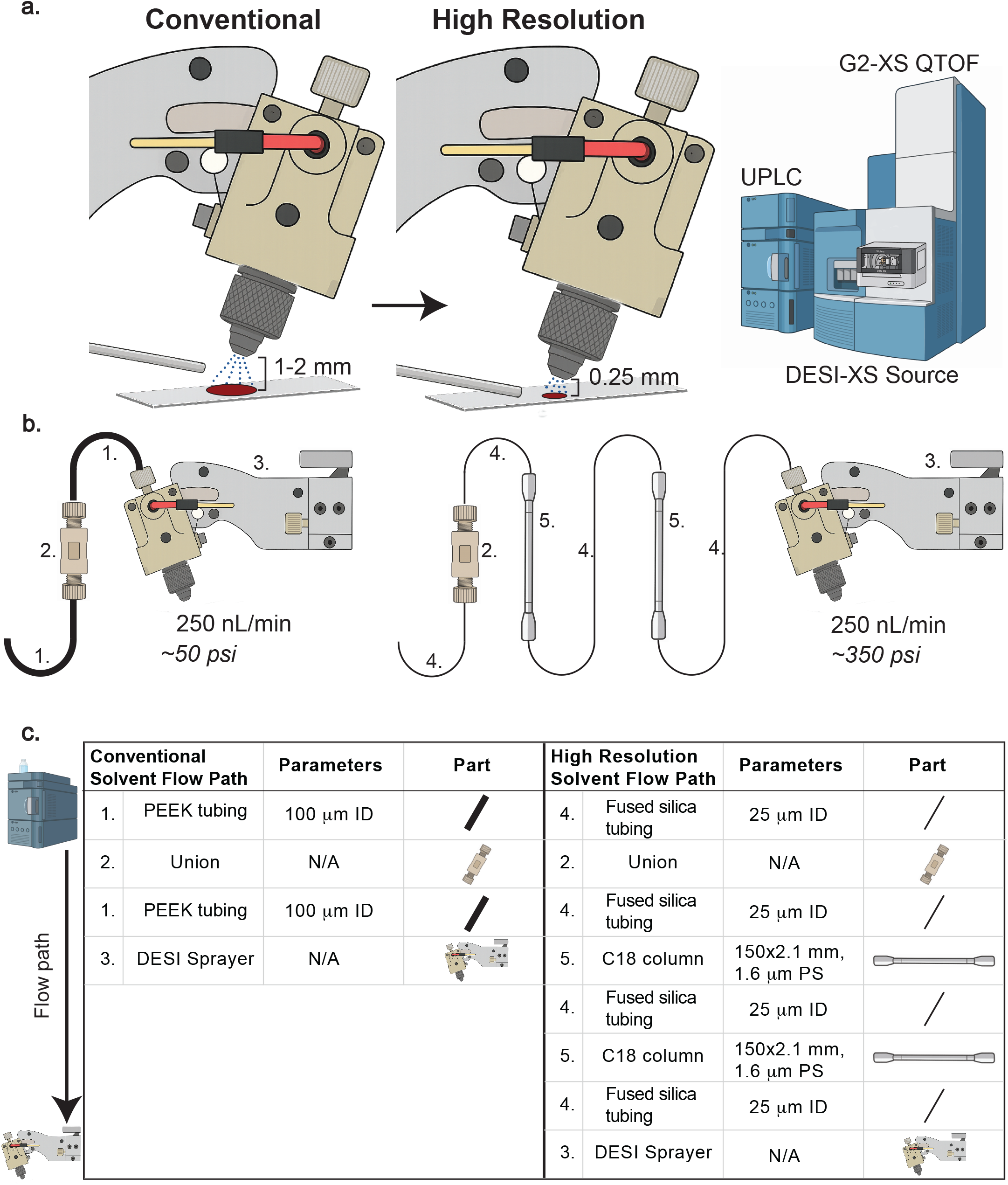
DESI-MSI hardware full diagram. **a,** Diagram showing the reduced emitter-to-sample distance. **b,** Diagram showing the hardware changes necessary for increased backpressure. c, List of solvent-flow-path components outlined in b.

**Supplementary Figure 2.**
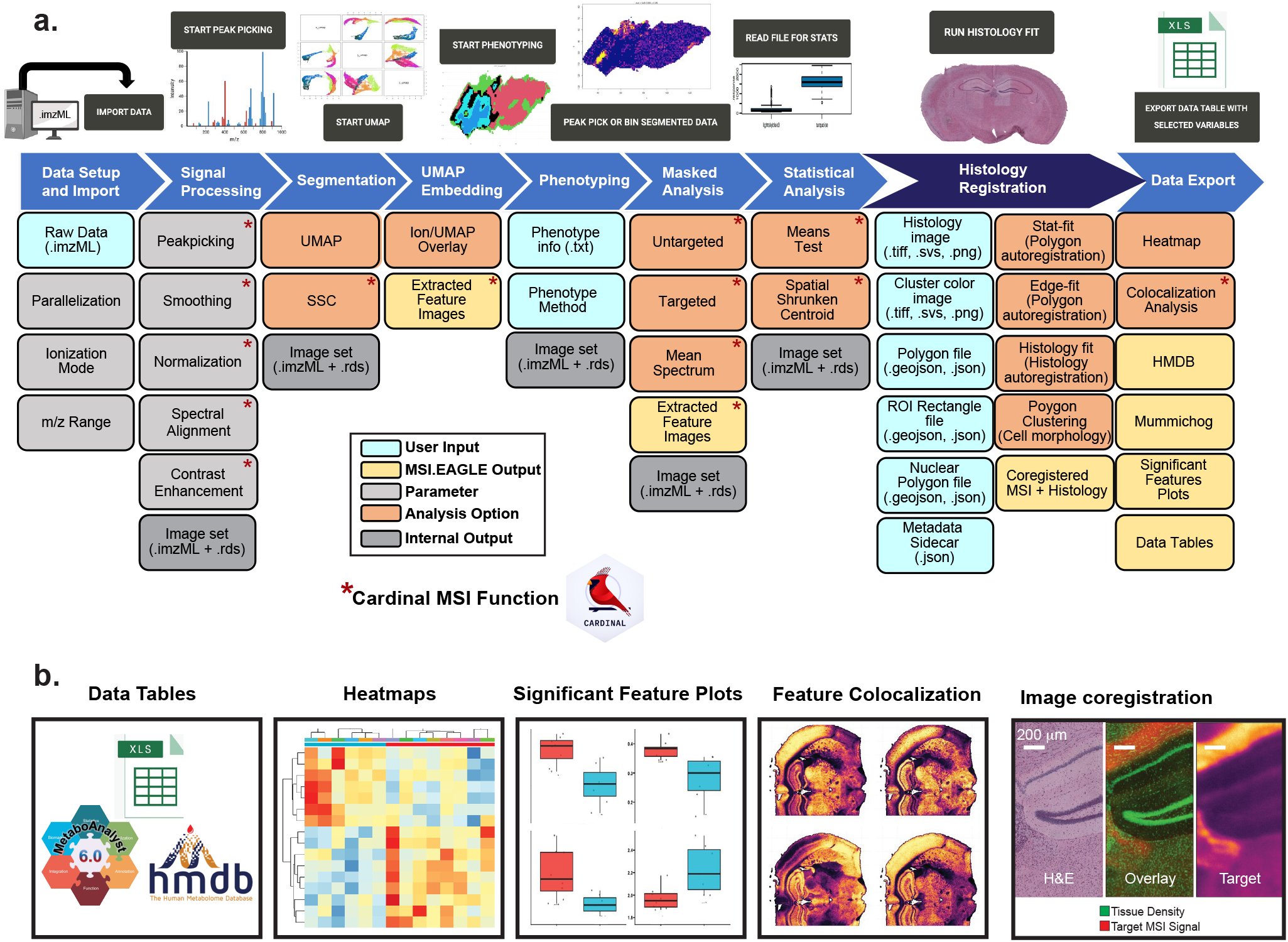
MSI.EAGLE pipeline. **a,** Analytical pipeline for processing and interpretation of mass spectrometry imaging (MSI) data. The workflow progresses through nine sequential modules: data import of raw imzML files, signal processing including peak picking and normalization, spatial segmentation, dimensionality reduction via UMAP embedding, phenotype definition, masking of regions of interest, statistical analysis, histology registration, and data export. Color-coding indicates functional classifications: user inputs (blue), MSI.EAGLE outputs (yellow), adjustable parameters (light gray), analysis options (orange), and internal outputs (dark gray). Interactive entry points specifying each module in the top section (black buttons) allow users to initiate key analytical processes. Red asterisks denote functions that leverage the Cardinal MSI package. **b,** Standardized outputs generated by MSI.EAGLE for downstream analysis and interpretation, including structured data tables for secondary analysis, interactive heatmaps for metabolite abundance visualization, significant feature plots highlighting statistical differences between regions, feature colocalization maps for spatial correlation analysis, and coregistration of histology data. The integrated platform facilitates cohesive progression from raw MSI data to biologically meaningful features through a unified interface, enabling systematic tissue characterization and biomarker discovery.

**Supplementary Figure 3.**
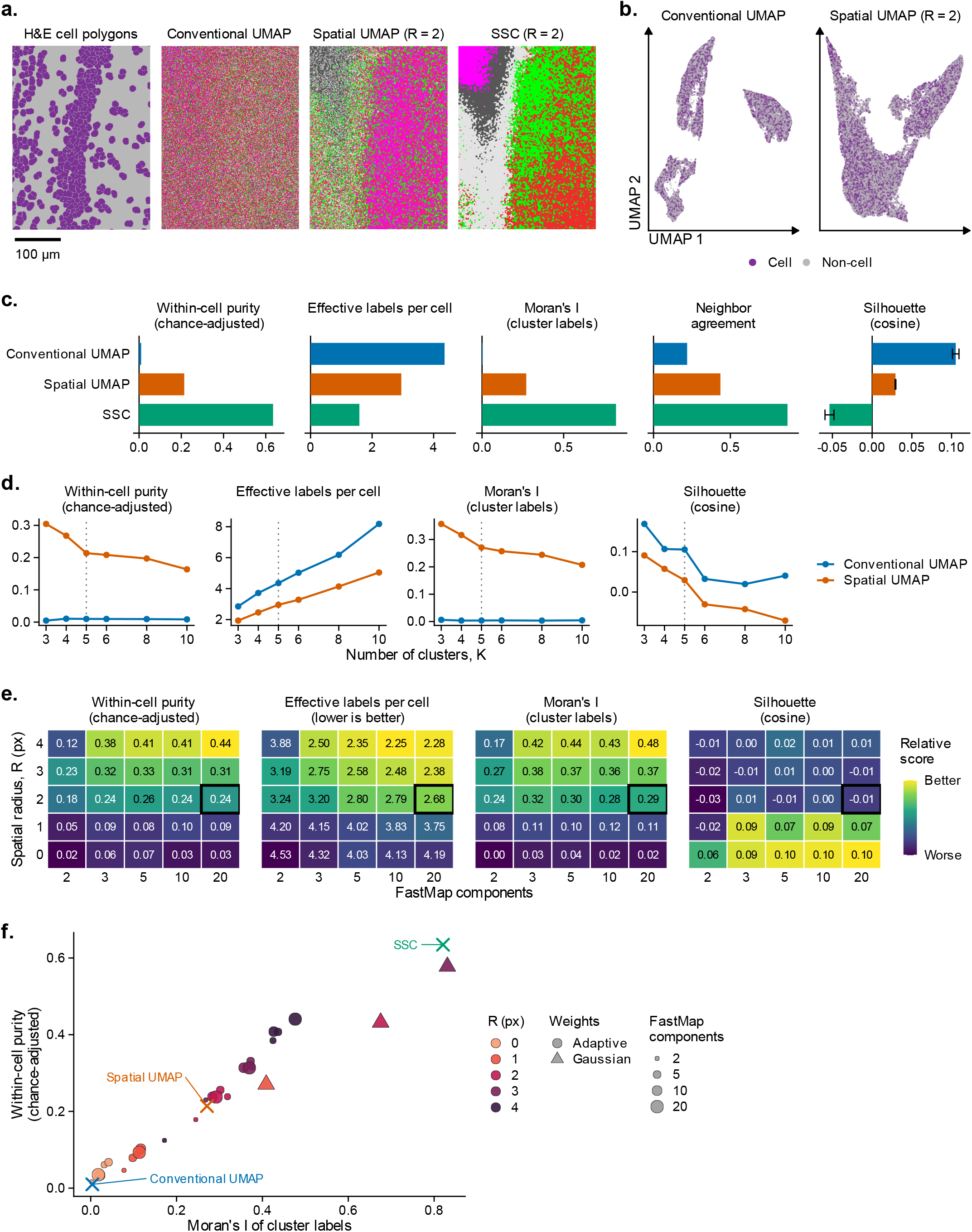
Quantitative benchmarking of spatial UMAP. **a**, Cluster maps over the full field (x = 401-700 µm, y = 101-500 µm) for the H&E cell reference (purple, cell; gray, non-cell), conventional UMAP, spatial UMAP (R = 2) and SSC (R = 2), each at K = 5. Cluster labels are arbitrary; for continuity with Fig. 3, each cluster is drawn in the color of the Fig. 3 cluster it overlaps most (optimal one-to-one matching; SSC clusters matched to those of spatial UMAP). Scale bar, 100 µm. **b**, First two dimensions of the conventional and spatial UMAP embeddings (40,000 randomly subsampled pixels), colored by whether each pixel falls inside a registered cell polygon (purple, cell; gray, non-cell). **c**, Five metrics for each method (blue, conventional UMAP; orange, spatial UMAP; green, SSC). The H&E reference segmentation is not plotted because its purity and labels per cell are 1.0 by construction (its Moran’s I is 0.856, neighbor agreement 0.931 and silhouette −0.0004). Within-cell purity (chance-adjusted): for each cell, the fraction of its pixels carrying that cell’s modal cluster label, averaged over cells, then corrected as (observed - chance)/(1 - chance), where chance is the same quantity recomputed after five random permutations of the cluster-label vector (identical permutations for every method); correction is necessary because raw purity is inflated by small K and by uneven cluster sizes. Effective labels per cell: inverse-Simpson count 1/Σp², where p is the fraction of a cell’s pixels in each cluster, averaged over cells; 1 denotes a perfectly intact cell and lower is better. Moran’s I: each cluster is converted to a 0/1 indicator, and Moran’s I is computed under row-standardized rook (4-neighbor) contiguity on the 1 µm pixel grid; the plotted value is the prevalence-weighted mean across clusters. Neighbor agreement: fraction of rook-adjacent pixel pairs assigned the same cluster label. Silhouette (cosine): mean silhouette width in the original 708-dimensional feature space under cosine distance, averaged over five random subsamples of 5,000 pixels (the same subsamples for every method); error bars are the s.d. across subsamples. **d**, The metrics in c recomputed for K = 3-10 by re-running k-means on the conventional and spatial UMAP embeddings, showing that the ranking does not depend on clustering granularity; dotted line, K = 5 as in c. **e**, Spatial UMAP parameter sensitivity: each cell is one (spatial radius R, FastMap component count) combination, re-embedded and re-clustered from the spectra and scored as in c. R = 0 gives each pixel a neighborhood of itself and is therefore the non-spatial control. Printed values are the actual metric; fill is rescaled independently within each panel, with yellow always the better-performing direction (inverted for effective labels per cell, where lower is better). The boxed cell (R = 2, 20 components) is the spatial UMAP setting used in a-d and Fig. 3. **f**, Spatial coherence (Moran’s I) against chance-adjusted within-cell purity for all 25 settings in e plus the purely spatial (gaussian) weighting variants; point fill denotes R, point size denotes FastMap component count and shape denotes weighting scheme. Crosses mark the three methods scored in c, colored as in c.

**Supplementary Figure 4.**
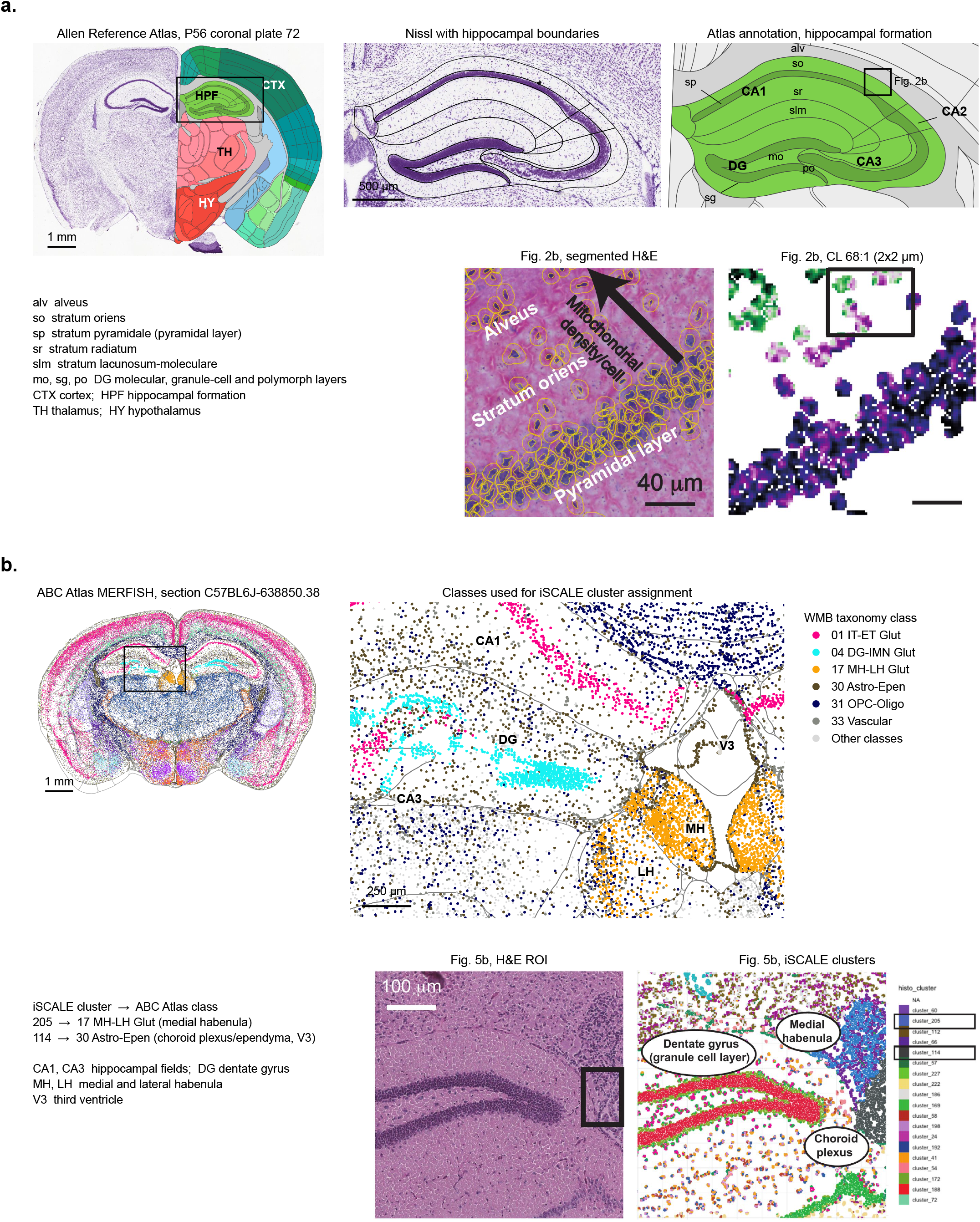
Allen Brain Atlas references. Allen Brain Atlas references used for anatomical (Fig. 2) and cell-type (Fig. 5) annotations. **a**, Hippocampal layer annotation for Fig. 2. Left, Allen Reference Atlas, adult (P56) mouse, coronal plate 72 of 132: the Nissl-stained reference section with the atlas structure annotation overlaid on the right hemisphere (CTX, cortex; HPF, hippocampal formation; TH, thalamus; HY, hypothalamus). The box marks the region enlarged at right. Middle, the Nissl section with the atlas hippocampal boundaries overlaid. Right, the atlas annotation of the same region, labeling fields CA1-CA3, the dentate gyrus (DG), the layers of CA1 (alv, alveus; so, stratum oriens; sp, stratum pyramidale; sr, stratum radiatum; slm, stratum lacunosum-moleculare) and the layers of DG (mo, molecular; sg, granule cell; po, polymorph). The box in dorsal CA1 marks the approximate position of the field imaged in Fig. 2b, reproduced below: segmented H&E with the alveus, stratum oriens and pyramidal layer annotated (arrow, increasing mitochondrial density per cell), and the matched 2×2 µm CL 68:1 ion image. Atlas panels were rendered from the vector annotation and Nissl image of the Allen Reference Atlas (atlas image 100960236), retrieved through the Allen Brain Map API. **b**, Cell-type reference for Fig. 5. Left, coronal MERFISH section C57BL6J-638850.38 from the Allen Brain Cell (ABC) Atlas whole-mouse-brain dataset, with each of its 120,186 cells plotted at its CCFv3-reconstructed position and colored by whole-mouse-brain taxonomy class; gray lines are CCFv3 region boundaries. The box marks the region enlarged at right, taken from the left hemisphere to match the orientation of Fig. 5. Right, the enlarged region with the six classes used for assignment shown in their ABC Atlas colors and all other classes in light gray: 01 IT-ET Glut (CA pyramidal neurons), 04 DG-IMN Glut (dentate granule cells), 17 MH-LH Glut (medial and lateral habenula), 30 Astro-Epen (astrocytes and the ependymal/choroid plexus lining of the third ventricle, V3), 31 OPC-Oligo and 33 Vascular. Below, the H&E ROI and iSCALE cluster map from Fig. 5b. Each iSCALE cluster was assigned the class whose spatial distribution it matched: cluster 205 to 17 MH-LH Glut and cluster 114 to 30 Astro-Epen (both boxed in the legend). Scale bars: a, 1 mm (overview), 500 µm (Nissl) and 40 µm (H&E); b, 1 mm (overview), 250 µm (enlargement) and 100 µm (H&E).

## Notes

https://github.com/Weljie-Lab-UPenn/MSI.EAGLE

https://zenodo.org/records/22943183

### Competing Interest Statement

The authors have declared no competing interest.

